# The *rhg1-a* (*Rhg1* low-copy) nematode resistance source harbors a copia-family retrotransposon within the *Rhg1-*encoded α-SNAP gene

**DOI:** 10.1101/653568

**Authors:** Adam M. Bayless, Ryan W. Zapotocny, Shaojie Han, Derrick J. Grunwald, Kaela K. Amundson, Andrew F. Bent

**Affiliations:** Department of Plant Pathology, University of Wisconsin – Madison, Madison, WI 53706 USA

**Keywords:** Plant disease resistance, retrotransposon, soybean cyst nematode, *Rhg1*

## Abstract

Soybean growers widely use the <u>R</u>esistance to *<u>H</u>eterodera <u>g</u>lycines* 1 (*Rhg1*) locus to reduce yield losses caused by soybean cyst nematode (SCN). *Rhg1* is a tandemly repeated four gene block. Two classes of SCN resistance-conferring *Rhg1* haplotypes are recognized: *rhg1-a* (“Peking-type”, low copy number, 3 or fewer *Rhg1* repeats) and *rhg1-b* (“PI 88788-type”, high copy number, 4 or more *Rhg1* repeats). The *rhg1-a* and *rhg1-b* haplotypes encode α-SNAP (alpha-<u>S</u>oluble <u>N</u>SF <u>A</u>ttachment <u>P</u>rotein) variants α-SNAP*_Rhg1_*LC and α-SNAP*_Rhg1_*HC respectively, with differing atypical C-terminal domains, that contribute to SCN-resistance. Here we report that *rhg1-a* soybean accessions harbor a copia retrotransposon within their *Rhg1 Glyma.18G022500* (α-SNAP-encoding) gene. We termed this retrotransposon “*RAC*”, for *<u>R</u>hg1* <u>a</u>lpha-SNAP <u>c</u>opia. Soybean carries multiple *RAC*-like retrotransposon sequences. The *Rhg1 RAC* insertion is in the *Glyma.18G022500* genes of all true *rhg1-a* haplotypes we tested and was not detected in any examined *rhg1-b* or *Rhg1_WT_* (single-copy) soybeans. *RAC* is an intact element residing within intron 1, anti-sense to the *rhg1-a α-SNAP* open reading frame. *RAC* has intrinsic promoter activities, but overt impacts of *RAC* on transgenic α-SNAP*_Rhg1_*LC mRNA and protein abundance were not detected. From the native *rhg1-a RAC^+^* genomic context, elevated α-SNAP*_Rhg1_*LC protein abundance was observed in syncytium cells, as was previously observed for α-SNAP*_Rhg1_*HC (whose *rhg1-b* does not carry *RAC*). Using a SoySNP50K SNP corresponding with *RAC* presence, just ∼42% of USDA accessions bearing previously identified *rhg1-a* SoySNP50K SNP signatures harbor the *RAC* insertion. Subsequent analysis of several of these putative *rhg1-a* accessions lacking *RAC* revealed that none encoded *α-SNAP_Rhg1_LC*, and thus they are not *rhg1-a*. *rhg1-a* haplotypes are of rising interest, with *Rhg4*, for combating SCN populations that exhibit increased virulence against the widely used *rhg1-b* resistance. The present study reveals another unexpected structural feature of many *Rhg1* loci, and a selectable feature that is predictive of *rhg1-a* haplotypes.

## Introduction

To thrive in their natural environments, organisms must continually sense and respond to changing conditions, including biotic and abiotic stresses. Transposable elements can cause relatively stable variation in numerous plant phenotypes such as flowering time, trichome presence or fruit size (Lisch, 2013). Transposable elements (TEs) may insertionally disrupt genes, or if TE activity is repressed by epigenetic transcriptional silencing, small interfering RNAs and chromatin condensation, this can impact the expression of nearby genes (Sigman and Slotkin, 2016). TEs are also increasingly being identified as modulatory factors during periods of host stress (i.e., heat, pathogen) (Liu et al., 2004; Xiao et al., 2008; Ding et al., 2015). For instance, cis-regulatory motifs within certain TEs can recruit stress-responsive host transcriptional factors, thereby influencing nearby host gene expression and potentially conferring a host benefit (Slotkin and Martienssen, 2007; Woodrow et al., 2010; Matsunaga et al., 2012; McCue and Slotkin, 2012; Cavrak et al., 2014; Makarevitch et al., 2015; Matsunaga et al., 2015; Negi et al., 2016; Galindo-Gonzalez et al., 2017). Some TEs also beneficially modulate the expression of host plant defense or susceptibility genes (Tsuchiya and Eulgem, 2013; Berg et al., 2015). Additionally, numerous studies report TEs lying within or adjacent to putative plant immune genes, however, potential influences on host genes or positive effects are often not apparent (Bhattacharyya et al., 1997; Henk et al., 1999; Wawrzynski et al., 2008).

*Glycine max* (soybean) is an important food and industrial crop (Schmutz et al., 2010). A major pest afflicting global soybean production is the soybean cyst nematode (SCN, *Heterodera glycines*), which causes yearly U.S. soybean yield losses of over 1 billion USD (Niblack et al., 2006; Mitchum, 2016; T. W. Allen, 2017). SCN is an obligate parasite that invades host roots and induces individual host cells to form a complex syncytium that serves as the SCN feeding site (Niblack et al., 2006; Mitchum, 2016). SCN feeding depletes available host resources and a functional syncytium must be maintained for 2-4 weeks for the nematode to complete its lifecycle. Since the unhatched eggs within cysts can remain viable for many years in the field, SCN is difficult to manage and is primarily controlled by growing naturally resistant soybeans (Niblack et al., 2006). Among known soybean loci contributing to SCN-resistance, the *Rhg1* (<u>R</u>esistance to *<u>H</u>eterodera <u>g</u>lycines* 1) locus found on chromosome 18 provides the strongest protection (Concibido et al., 2004). *Rhg1* causes the SCN-induced syncytium to fail a few days after induction, and the soybean PI 88788-type “*rhg1-b*” haplotype is the primary SCN resistance locus used in commercially grown soybeans (Concibido et al., 2004; Niblack et al., 2006; Mitchum, 2016).

Soybean *Rhg1* is an unusual disease resistance locus that consists of a ∼31.2 kb unit that is tandemly repeated as many as 10 times (Cook et al., 2012). Within each 31.2 kb *Rhg1* repeat unit are four different *Rhg1*-encoded genes: *Glyma.18G022400*, *Glyma.18G022500*, *Glyma.18G022600*, and *Glyma.18G022700*, none of which have similarity to previously identified resistance genes (Cook et al., 2012; Cook et al., 2014; Lee et al., 2015). Of the three *Rhg1* genes that contribute to SCN-resistance, only *Glyma.18G022500*, an α-SNAP (alpha-<u>S</u>oluble <u>N</u>SF <u>A</u>ttachment <u>P</u>rotein), has amino acid polymorphisms relative to the wild-type *Rhg1* gene alleles present in SCN-susceptible soybeans (Cook et al., 2012; Cook et al., 2014; Lee et al., 2015). The mRNA transcript abundance of all three resistance-associated *Rhg1* genes is significantly elevated in SCN-resistant multi-copy *Rhg1* soybeans, relative to SCN-susceptible single-copy *Rhg1* (WT *Rhg1*) soybeans (Cook et al., 2012; Cook et al., 2014).

At least two distinct *Rhg1* genotype classes exist: “low-copy *Rhg1*” (*rhg1-a*, sometimes referred to as *Rhg1_LC_*, often derived from PI 548402 ‘Peking’), and “high-copy *Rhg1*” (*rhg1-b*, sometimes referred to as *Rhg1_HC_*, often derived from PI 88788) (Niblack et al., 2002; Brucker et al., 2005; Cook et al., 2012; Cook et al., 2014; Bayless et al., 2018). These *Rhg1* genotype classes represent two distinct multi-copy *Rhg1* haplotypes that vary most notably by: a) *Rhg1* repeat number (a high or low number of *Rhg1* repeats), and b) encoding distinctive resistance-type α-SNAP proteins with C-terminal polymorphisms at a conserved functional site (Cook et al., 2014; Bayless et al., 2016). *rhg1-a* resistance is bolstered by an unlinked chromosome 8 locus, *Rhg4*, whose presence contributes to full-strength “Peking-type” SCN resistance (Meksem et al., 2001; Liu et al., 2012). *Rhg4* encodes a polymorphic serine hydroxy methyl transferase with altered enzyme kinetics, but the molecular basis of resistance augmentation by *Rhg4* is not yet understood (Liu et al., 2012; Mitchum, 2016). Several *rhg1-b* and *rhg1-a* accessions have been analyzed by whole genome sequencing (WGS) studies, and characteristic single nucleotide polymorphism (SNP) signatures predictive of *rhg1-b* or *rhg1-a* haplotype soybeans have been reported (Cook et al., 2014; Lee et al., 2015; Shi et al., 2015; Kadam et al., 2016; Patil et al., 2019). Additionally, studies by Arelli, Young and others have profiled SCN-resistance among thousands of USDA soybean accessions and noted substantial phenotypic variation (e.g., (Anand, 1984; Hussey et al., 1991; Young, 1995; Diers et al., 1997; Arelli et al., 2000; Vuong et al., 2015; Klepadlo et al., 2018)). However, the influence of all *Rhg1* haplotype and/or allelic variation factors on SCN resistance expression or plant yield is not yet fully understood.

Several recent studies have deepened our understanding of *Rhg1* molecular function and highlight a central role of the SNARE (<u>S</u>oluble <u>N</u>SF <u>A</u>ttachment Protein <u>RE</u>ceptors)-recycling machinery in SCN-resistance (Matsye et al., 2012; Cook et al., 2014; Bayless et al., 2016; Lakhssassi et al., 2017; Bayless et al., 2018). α-SNAP, and the ATPase NSF (<u>N</u>-ethylmaleimide <u>S</u>ensitive <u>F</u>actor), are conserved eukaryotic housekeeping proteins that form the core SNARE-recycling machinery. They sustain the pool of fusion-competent SNAREs necessary for new membrane fusion events (Sudhof and Rothman, 2009; Zhao et al., 2015). While most animals encode single NSF and α-SNAP genes, soybean is a paleopolyploid that encodes two NSF, four or five α-SNAP and two γ-SNAP (gamma-SNAP) genes, respectively. A C-terminal α-SNAP domain conserved across all plants and animals recruits NSF to SNARE-bundles and stimulates the ATPase activity of NSF that powers SNARE-complex recycling. However, it is this otherwise conserved α-SNAP C-terminal region that is atypical among both *rhg1-b*- and *rhg1-a-* encoded α-SNAP proteins, and accordingly, both *Rhg1* resistance type α-SNAPs are impaired in promoting normal NSF function and instead mediate dosage-dependent cytotoxicity (Sudhof and Rothman, 2009; Cook et al., 2014; Zhao et al., 2015; Bayless et al., 2016). The abundance of the atypical *rhg1-b* α-SNAP*_Rhg1_*HC protein specifically increases in the SCN feeding site and contributes to *Rhg1-*mediated collapse of the SCN-induced syncytium (Bayless et al., 2016).

At least two additional loci associated with SCN-resistance are also components of the SNARE-recycling machinery (Lakhssassi et al., 2017; Bayless et al., 2018). Recently, a specialized allele of NSF, *NSF_RAN07_* (*<u>R</u>hg1-*<u>a</u>ssociated <u>N</u>SF on chromosome <u>07</u>), was shown to be necessary for the viability of *Rhg1-*containing soybeans (Bayless et al., 2018). Compared to the WT NSF_Ch07_ protein, the NSF_RAN07_ protein more effectively binds to resistance type α-SNAPs and confers better protection against resistance-type α-SNAP-induced cytotoxicity (Bayless et al., 2018). During the *Rhg1-*mediated resistance response, the ratio of *Rhg1* resistance-type to WT α-SNAPs increases and is apparently an important factor underlying resistance (Bayless et al., 2016; Bayless et al., 2018). Two genetic events sharply reduce WT α-SNAP protein abundance in SCN-resistant *rhg1-a* soybeans (Bayless et al., 2018). First, the wild type α-SNAP-encoding block at *Rhg1* on chromosome 18 – a predominant source of total WT α-SNAP proteins in soybean – is absent from all examined *rhg1-a* accessions, thereby diminishing overall WT α-SNAP protein abundance (Cook et al., 2014; Bayless et al., 2018). Secondly, *rhg1-a* lines often carry a null allele of the α-SNAP encoded on chromosome 11 (*Glyma.11G234500*) – the other major source of WT α-SNAP proteins – due to an intronic splice site mutation that causes premature translational termination and loss of protein stability (Matsye et al., 2012; Cook et al., 2014; Lakhssassi et al., 2017; Bayless et al., 2018). Together, the above studies support a paradigm whereby *Rhg1* and associated SCN-resistance loci rewire major components of the soybean SNARE-recycling machinery. Importantly, soybean accessions that carry *rhg1-a* and *Rhg4* can resist many of the virulent SCN populations that partially overcome *rhg1-b* resistance (Brucker et al., 2005; Niblack et al., 2008; Bayless et al., 2016). Therefore, there is considerable interest in understanding and using *rhg1-a*, the subject of the present study, as an alternative to *rhg1-b* in commercial soybean cultivars (Brucker et al., 2005; Liu et al., 2012; Yu et al., 2016).

Presence/absence variation of transposable elements at specific loci is common among different soybean accessions, and tens of thousands of non-reference genome TE insertions occur between cultivated and wild soybean (Tian et al., 2012). Moreover, high TE densities near genomic regions exhibiting structural polymorphisms such as copy number variation are also reported in soybean (McHale et al., 2012). While examining the *Rhg1* low-copy (*rhg1-a*) haplotype of soybean accession PI 89772, we uncovered an intact copia retrotransposon within all three copies of the *Rhg1*-encoded *α-SNAP* genes. We termed this retrotransposon “*RAC*”, for *<u>R</u>hg1* <u>a</u>lpha-SNAP <u>c</u>opia). The *RAC* element, which is entirely within the first intron of the *Glyma.18G022500* (α-SNAP) gene, appears to be intact and transcribes anti-sense to the *α-SNAP* ORF. BLAST searches revealed similar copia elements across the soybean genome, suggesting why assemblies of Illumina short-read whole-genome sequences failed to include this sequence within *rhg1-a* assemblies. This *α-SNAP-RAC* insertion was absent from all examined single-copy *Rhg1* (SCN-susceptible) and high-copy *rhg1-b* (*Rhg1_HC_*) accessions. More than half of the USDA accessions with SoySNP50K SNPs preliminarily indicative of a low-copy *rhg1-a* haplotype did not carry *RAC*, but sub-sampling among those accessions revealed that they do not encode α-SNAP*_Rhg1_*LC and thus are not *rhg1-a*. The increasingly important *rhg1-a* SCN-resistant soybean breeding lines do harbor this previously unreported retrotransposon within the α-SNAP*_Rhg1_*LC-encoding gene.

## Materials & Methods

### Transgenic soybean hairy root generation and root culturing

Transgenic soybean roots were produced using *Agrobacterium rhizogenes* strain “ARqua1” (Quandt et al., 1993) and the previously described binary vector pSM101, as in (Cook et al., 2012). Transgenic roots were sub-cultured in the dark at room temperature on hairy root medium as in (Cook et al., 2012).

### DNA extraction

Soybean genomic DNAs were extracted from expanding trifoliates or root tissues of the respective soybean accessions using standard CTAB methods similar to (Cook et al., 2012).

### Amplification and detection of *RAC* (*Rhg1* alpha-SNAP copia)

For initial amplification and subcloning of native *α-SNAP-RAC*, approximately 100 ng of CTAB-extracted gDNA from PI 89772 (*rhg1-a*) was PCR amplified for 35 cycles using HiFi polymerase (KAPA Biosystems, Wilmington, MA). Primer annealing was at ∼70°C for 30 seconds and extension was at 72°C for 5 minutes. The resulting *α-SNAP-RAC* amplicon from PI 89772 was separated by agarose gel electrophoresis, gel extracted using a Zymoclean Large Fragment DNA Recovery Kit (Zymo Research, Irvine CA) and TA overhang cloned into a pTopo xL vector using the Topo xL PCR Cloning Kit (Life Technologies Corp., Carlsbad CA), per manufacturer’s recommendations. For PCR detection of *α-SNAP-RAC* junctions or WT exon distances, ∼25 ng of CTAB-extracted genomic DNA from each respective accession was amplified using GoTAQ Green (New England Biolabs, Ipswich MA) for 32 cycles, separated on a 0.8% agarose gel and visualized.

### Phylogenetic tree construction

For the *RAC-*like nucleotide tree, evolutionary analyses were conducted in MEGA7 (Kumar et al., 2016) and evolutionary history was inferred by using the Maximum Likelihood method based on the Tamura-Nei model (Tamura et al., 2004). The tree with the highest log likelihood (−47751.11) is shown. Initial tree(s) for the heuristic search were obtained automatically by applying Neighbor-Join and BioNJ algorithms to a matrix of pairwise distances estimated using the Maximum Composite Likelihood (MCL) approach, and then selecting the topology with superior log likelihood value. The analysis involved 24 nucleotide sequences. All positions containing gaps and missing data were eliminated. There were a total of 4157 positions in the final dataset.

For the *RAC-*polyprotein tree, evolutionary analyses were conducted in MEGA7 (Kumar et al., 2016) and the evolutionary history was inferred by using the Maximum Likelihood method based on the JTT matrix-based model (Jones et al., 1992). The tree with the highest log likelihood (−12400.87) is shown. Initial tree(s) for the heuristic search were obtained automatically by applying Neighbor-Join and BioNJ algorithms to a matrix of pairwise distances estimated using a JTT model, and then selecting the topology with superior log likelihood value. The tree is drawn to scale, with branch lengths measured in the number of substitutions per site. The analysis involved 14 amino acid sequences. All positions containing gaps and missing data were eliminated. There were a total of 644 positions in the final dataset.

### Read Depth Analysis of *RAC*

Using previously published whole-genome-sequencing data (Cook et al., 2014), read depth was computed using the depth program of SAMtools (Li et al., 2009). Depth was averaged in 250 bp intervals on Chromosome 10 from bp 40,650,000 – 40,690,000 (includes flanking regions of the 99.7% identity *RAC*-like element). The copy number of the Chromosome 10 *RAC*-like element was then calculated as the ratio of the read coverage per 250 bp from bp 40,672,000 – 40,675,750, divided by the average read coverage for the flanking regions between bp 40,650,000 – 40,690,000. Sequencing coverage was visualized using ggplot (Wickham, 2009) within RStudio (RStudio Team, 2015).

### Methylation analysis

McrBC methylation studies were performed similarly to (Cook et al., 2014). Control McrBC reactions contained equivalent amounts of gDNA in reaction buffer, but had no added McrBC enzyme. McrBC digestion was performed at 37°C for 90 minutes, followed by a 20 minute heat inactivation at 65°C. McrBC digested or mock treated samples were PCR amplified with primers flanking 5’ or 3’ *α-SNAP-RAC* junctions and visualized by agarose gel electrophoresis.

### RNA isolation and cDNA synthesis

Total RNAs were extracted using Trizol (Life Technologies Corp., Carlsbad CA) or the Direct-Zol RNA Miniprep Kit (Zymo Research, Irvine CA), per manufacturer’s instructions. All RNA samples were DNAase treated and quantified using a spectrophotometer. cDNA synthesis was performed using the iScript cDNA Synthesis Kit (Bio-Rad, Hercules CA) according to manufacturer’s recommendations using 1.0 µg of purified total RNA.

### qPCR analysis

qPCR was performed with a CFX96 real-time PCR detection system (BioRad Laboratories, Hercules CA) using SsoFast EvaGreen Supermix (Bio-Rad Laboratories, Hercules CA) as in (Cook et al., 2012). Following amplification, a standardized melting curve analysis program was performed. Overall cDNA abundances for each sample were normalized using the qPCR signal for reference gene *Glyma.18G022300*. *RAC* transcript abundances are presented relative to the mean abundance of *RAC* transcript for Williams 82 leaf samples.

### RT-PCR analysis

For RT-PCR, 31 cycles of amplifications were performed prior to loading PCR product samples for separation and visualization by agarose gel electrophoresis. The number of PCR cycles terminated prior to maximal amplification of product from the most abundant template pool. A primer set sitting on a conserved copia region detected both endogenous *RAC-*like transcripts as well as the uniquely tagged *RAC* transgene. Specific detection of the tagged *RAC* transgene was with a primer pair sitting atop the engineered region. Transcript from *Glyma.18G022300* or *Skp16* served as a control for both cDNA quality and relative transcript abundance.

### Vector construction

Native *α-SNAP-RAC* was PCR amplified from pTopo XL subclones using Kapa HiFi polymerase with AvrII and SbfI restriction site overhangs. Following agarose gel (cut with XbaI/PstI) (New England Biolabs, Ipswich MA). Gel extractions were performed using the QIAquick Gel Extraction Kit (Qiagen, Hilden Germany) or the Zymoclean Large Fragment DNA Recovery Kit (Zymo Research, Irvine CA). Purified DNA fragments were ligated overnight at 4°C with T4 DNA ligase (New England Biolabs, Ipswich MA) per manufacturer’s recommendations. To remove *RAC* from within the native *α-SNAP-RAC* subclone in vector pSM101, the Polymerase Incomplete Primer Extension (PIPE) PCR method was used with Kapa HiFi polymerase (Klock and Lesley, 2009). Similarly, the synonymous tag added within the *RAC* ORF of native *α-SNAP-RAC* was created using PIPE. Unique nucleotide tag was located ∼160 bp downstream of the *RAC* ATG and maintains an intact ORF. For creating the *RAC* only vector which assessed inherent *RAC* transcription, the native *RAC* ORF with both LTRs (∼4.77 kb) was amplified from the initial PI 89772 *α-SNAP-RAC* subclone in pTopoXL using Kapa HiFi. Restriction site overhangs for AvrII and SbfI were incorporated into the primer sequences, and following gel recovery, the PCR amplicon was restriction digested and ligated into a PstI/XbaI cut pSM101 binary vector using T4 DNA ligase (New England Biolabs, Ipswich MA). For the native *α-SNAP-RAC* with flanking *Rhg1* sequence, an 11.1 kb native *Rhg1* sequence containing *Glyma.18G022400* and *Glyma.18G022500* (and ∼1 kb downstream of each stop codon), was PCR amplified from a previously published fosmid subclone “Fos-32”, with AvrII and SbfI restriction ends using Kapa HiFi polymerase (Cook et al., 2012). After restriction digestion, this amplified native fragment was ligated into the binary vector pSM101 (digested with PstI and XbaI). This created a native *Rhg1* two gene vector of the *rhg1-b* type. Then, to make the native *rhg1-a* construct, this native *rhg1-b* pSM101 vector was used as a scaffold for step-wise cloning of two different native *rhg1-a* fragments amplified from PI 89772 genomic DNA. The first was a 4 kb fragment with an SbfI primer overhang containing *Glyma.18G022400* up til an endogenous NruI site at exon 1; the second 11 kb fragment resumed at NruI until ∼ 1.0 kb downstream of the *Glyma.18G022500* (*α-SNAP_Rhg1_LC*) termination codon and contained an AvrII restriction overhang.

### Immunobloting & antibodies

Affinity-purified polyclonal rabbit antibodies raised against α-SNAP*_Rhg1_*LC and wild-type α-SNAP C-terminus were previously generated and validated in 1(Bayless et al., 2016). Protein lysates were prepared from ∼100 mg of soybean roots that were immediately flash-frozen in liquid N_2_. Roots were homogenized in a PowerLyzer 24 (Qiagen) for three cycles of 15 seconds, with flash-freezing in-between each cycle. Protein extraction buffer [50 mM Tris·HCl (pH 7.5), 150 mM NaCl, 5 mM EDTA, 0.2% Triton X-100, 10% (vol/vol) glycerol, 1/100 Sigma protease inhibitor cocktail] was then added at a 3:1 volume to mass ratio. Lysates were then centrifuged at 10,000g for 10 minutes and supernatant was added to SDS-PAGE loading buffer. Immunoblots were performed essentially as in (Song et al., 2015; Bayless et al., 2016). Briefly, immunoblots for α-SNAP*_Rhg1_*LC or WT α-SNAPs were incubated overnight at 4°C in 5% (wt/vol) nonfat dry milk TBS-T (50 mM Tris, 150 mM NaCl, 0.05% Tween 20) at 1:1,000. NSF immunoblots were performed similarly. Secondary horseradish peroxidase-conjugated goat anti-rabbit IgG was added at 1:10,000 and incubated for 1 h at room temperature on a platform shaker, followed by four washes with TBS-T. Chemiluminescence signal detection was performed with SuperSignal West Pico or Dura chemiluminescent substrates (Thermo Scientific, Waltham WA) and developed using a ChemiDoc MP chemiluminescent imager (Bio-Rad, Hercules CA).

### Immunolabeling and Electron Microscopy

Immunolabeling was performed as in (Bayless et al., 2016). Transverse sections of ∼2 mm long soybean (cv. Forrest) root areas containing syncytia were harvested by hand-sectioning at 4 dpi. Root sections were fixed in 0.1% glutaraldehyde and 4% (vol/vol) paraformaldehyde in 0.1M sodium phosphate buffer (PB) (pH 7.4) overnight after vacuum infiltration for about 1 hour. After dehydration in ethanol, samples were then embedded in LR White Resin. Ultrathin sections (∼90-nm) were taken longitudinally along the embedded root pieces using an ultramicrotome (UC-6; Leica) and mounted on nickel slot grids. For the immunogold labeling procedure, grids were first incubated on drops of 50 mM glycine/PBS for 15 min, and then blocked in drops of blocking solutions for goat gold conjugates (Aurion; Wageningen, NL) for 30 min and then equilibrated in 0.1% BSA-C/PBS (incubation buffer). Grids were then incubated overnight at 4°C with custom *α*-SNAP*_Rhg1_*LC polyclonal antibody (diluted 1:1000 in incubation buffer), washed five times in incubation buffer, and incubated for 2 h with goat anti-rabbit antibody conjugated to 15-nm gold (Aurion) diluted 1:50 in incubation buffer. After six washes in incubation buffer and two 5-min washes in PBS, the grids were fixed for 5 min in 2.0% (vol/vol) glutaraldehyde in 0.1 M phosphate buffer, followed by two 5-min washes in 0.1 M phosphate buffer and five 2-min washes in water. Images were collected with a MegaView III digital camera on a Philips CM120 transmission electron microscope.

### Oligonucleotide Primers

RAC qRT For: GGGTTCGAAATGAATACCTG

RAC qRT Rev: CACGTTCTTCTCATGGATCCTA

RAC Delete PIPE For: CTT CAT CCA CAA TTC TAA TTT ATA TGC TAG

RAC Delete PIPE Rev: GAA TTG TGG ATG AAG TAC GAC AAT CAA C

5’ RAC Junction For: TGGCTCCAAGTATGAAGATGCC

5’ RAC Junction Rev: AACTACAGTGGCTGACCTTCT

3’ RAC Junction For: ACTGTTCATTCAGACCGCGT

3’ RAC Junction Rev: GCAATGTGCAGCATCGACATGGG

WT Junction For: GAGTTTTGAGGTGTCCGATTTCCC WT

Junction Rev: GTGAGCGCAGTCACAAACAAC

5’ Methylation For: TGGCTCCAAGTATGAAGATGCC

5’ Methylation Rev: AACTACAGTGGCTGACCTTCT

3’ Methylation For: ACTGTTCATTCAGACCGCGT

3’ Methylation Rev: GCAATGTGCAGCATCGACATGGG

Skp16 qRT For: GAG CCC AAG ACA TTG CGA GAG

Skp16 qRT Rev: CGG AAG CGG AAG AAC TGA ACC

2570 qRT For: TGA GAT GGG TGG AGC TCA AGA AC

2570 qRT Rev: AGC TTC ATC TGA TTG TGA CAG TGC

For RAC Tag Mut: GCTCTGCTCCTGAGCCCTTGAAAACGGACAGAATGCACGGAG

Rev RAC Tag Mut: GCTCAGGAGCAGAGCCATCTATGAACTCCACTTTATTCTTGGC

RAC Tag Detect For: CAGTCCTAGACTCAACCAATTACC

RAC Tag Detect Rev: CCTTGGCTATACCTGCTCTTTAAATC

For RAC initial TopoXL subclone: GAGATTACATTGGATGATACGGTCGACC

Rev RAC initial TopoXL subclone: AGATAAGATCAGACTCCAGCAACCTC

For RAC Alone subclone AvrII: cctaggGGTGTCCGATTTCCCGATTAATTGAAG

Rev RAC Alone subclone SbfI: cctgcaggCCAACATCAATTTCAAAGTTCGTCACTTTC

LC-Splice Reverse: AGTAATAACCTCATACTCCTCAAGTT

LC-Splice Full For: GAGGAGGTTGTTGCTATAACCAATGC

LC-Splice Isoform For: GAGGAGGAACTGGATCCAACATTTTC

SbfI Native *Glyma.18G022400* For: cctgcaggGAGCAGTAGGCTTCTTTGGAACTTG

AvrII Native *Glyma.18g022500* Rev:cctaggGTTCCTAAAGTGGCAAACCCTAAGAACAAAG

BglII Native For: AGATCTCCCTGAGAGTATCTTGATTTCAGATCG

BglII Native Rev: AGATCTTTTACGCATATCCGACCTTCAAC

### Accession Numbers

NCBI Accessions will be established for the RAC sequences shown in Supplemental Figs S2 and S3.

### Supplemental Data files

*See: Bayless2019-SI-combined.pdf*

**Fig. S1.** Native genomic *rhg1-a* α-SNAP amplicons from PI 89772 are unexpectedly large and contain an inserted DNA element.

**Fig. S2.** Complete nucleotide sequence of the PI 89772 (*rhg1-a*, *Rhg1* low-copy) *RAC* (element and flanking exonic α-SNAP_Rhg1_LC regions.

**Fig. S3.** Translation of the *RAC*-encoded polyprotein (1438 residues) from accession PI 89772.

**Fig. S4.** Depth of short genomic reads that align to the *RAC* nucleotide sequence.

**Fig. S5.** Sequence of the *RAC* SNP (ss715606985), nucleotide SNP signatures of consensus *rhg1-a* (*Rhg1* low-copy) and *rhg1-b* (*Rhg1* high-copy) haplotypes, and detected *Rhg1* α-SNAP transcripts from RAC^+^ or RAC^−^ accessions with consensus *rhg1-a* SNP-signatures.

**Fig. S6.** Flow chart summarizing findings regarding RAC^+^ vs. RAC^−^ accessions, which otherwise have consensus SNP signatures for *rhg1-a*.

**Fig. S7.** α-SNAP_Rhg1_LC immunolabeling in ‘Forrest’ roots is highly specific.

**Table S1.** RAC-like elements identified from NBLAST searches of RAC against the Williams 82 reference genome at Phytozome.org.

## Results

### Multiple *rhg1-a* haplotypes harbor an intronic copia retrotransposon (*RAC*) within the ***Rhg1-*encoded *α-SNAP***

The α-SNAPs encoded by the *rhg1-a* and *rhg1-b* loci play a key role in SCN-resistance (Cook et al., 2012; Cook et al., 2014; Bayless et al., 2016; Liu et al., 2017; Bayless et al., 2018). While entire 31.2 kb *Rhg1* repeats of *rhg1-b* and *Rhg1_WT_*-like haplotypes have been sub-cloned and characterized (Cook et al., 2012), we sought to study the native genomic *rhg1-a* α-SNAP-encoding region to investigate potential regulatory differences between *rhg1-a* and *rhg1-b*. Fig 1A provides a schematic of the 31.2 kb *Rhg1* repeat unit with the four *Rhg1*-encoded genes, while Fig 1B presents a schematic of the previously published *Rhg1* haplotypes (single-, low-, and high-copy) and the respective C-terminal amino acid polymorphisms of their *Rhg1-*encoded α-SNAP proteins (Cook et al., 2012; Cook et al., 2014). Working from previously generated WGS data, we PCR-amplified the native genomic *rhg1-a α-SNAP* locus from PI 89772, and unexpectedly, obtained PCR amplicons ∼5 kb larger than WGS-based estimates (Fig S1A) (Cook et al., 2014; Liu et al., 2017). Overly large amplicons were also obtained using different PI 89772 genomic DNA templates and/or other PCR cycling conditions, and no *rhg1-a α-SNAP* amplicons of expected size were observed (Fig S1A). Sanger DNA sequencing of these unusually sized *rhg1-a α-SNAP* amplicons matched WGS predictions until part way into *α-SNAP* intron 1, where a 4.77 kb element was inserted. Immediately following this 4.77 kb insertion, the amplicon sequence again matched WGS predictions (Fig 1C).

**Fig 1.**
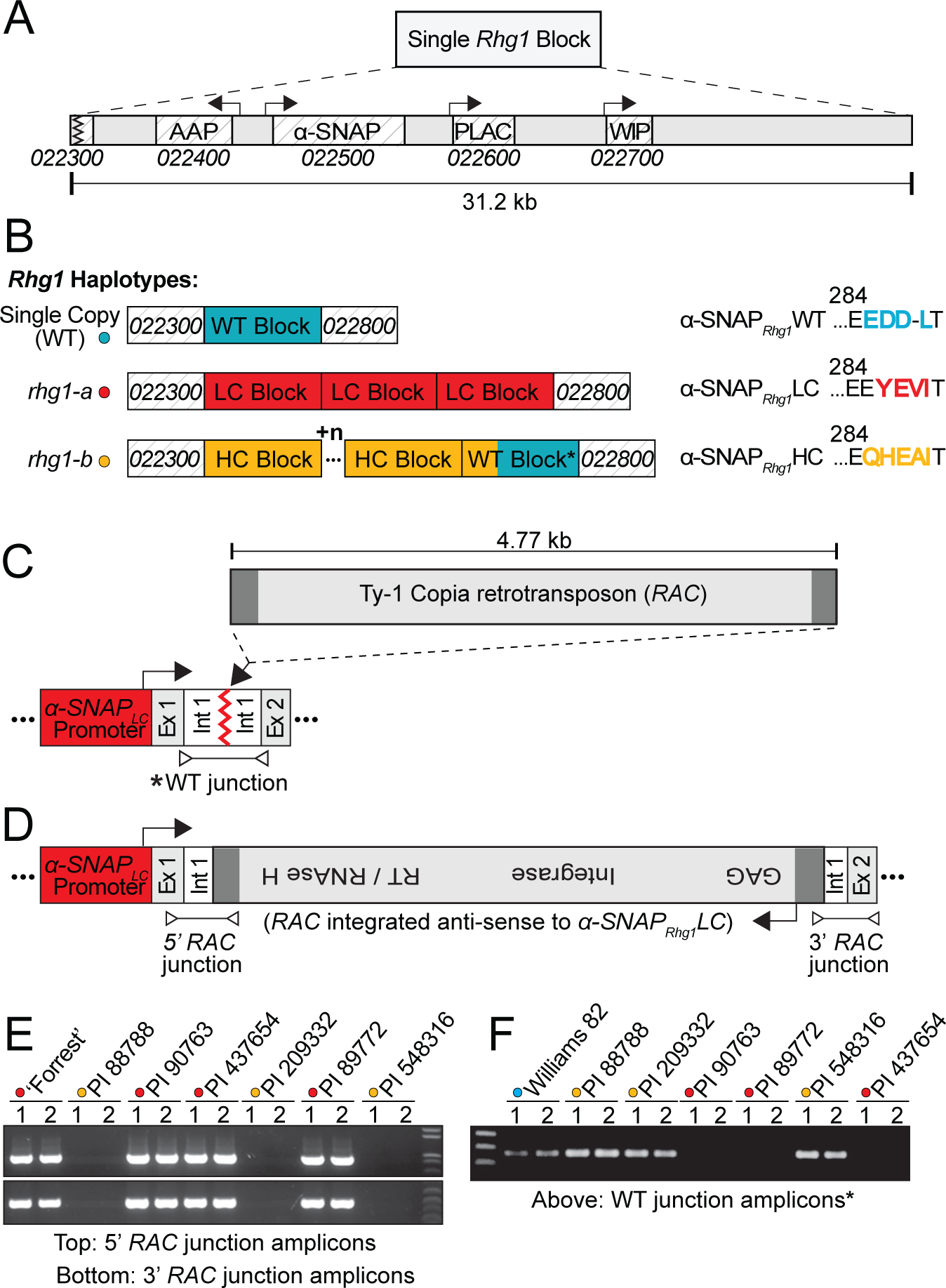
Multiple *rhg1-a* haplotypes harbor an intronic copia retrotransposon (*RAC*) within the *Rhg1-*encoded *α-SNAP* (*Glyma.18g022500*). (A) Diagram of a single 31.2 kb *Rhg1* block and the four *Rhg1*-encoded genes: *Glyma.18G022400* (amino acid permease, AAP), *Glyma.18G022500* (α-SNAP), *Glyma.18G022600* (PLAC-domain protein), and *Glyma.18G022700* (wound-inducible protein, WIP). *Glyma.18G022300* and *Glyma.18G022800* flank *Rhg1*, but each repeat also includes a truncated 3’ fragment of *Glyma.18G022300*. (B) Schematic of the three known *Rhg1* haplotypes: *Rhg1* wild-type (single-copy, shown blue), *rhg1-a* (low-copy, shown red) and *rhg1-b* (high-copy, shown orange). *Rhg1* α-SNAP C-terminal amino acid polymorphisms colored to match *Rhg1* block type. (C, D) Model from DNA sequencing of *Rhg1* alpha-SNAP copia (*RAC*) integration site within the PI 89772 (*rhg1-a*) encoded *α-SNAP*. The 4.77 kb *RAC* element (shown grey) is anti-sense to *α-SNAP_Rhg1_LC* and increases overall repeat size to ∼36 kb. *RAC* ORF is intact and encodes a 1438 residue polyprotein. *RAC* LTRs shown dark grey, *α-SNAP_Rhg1_LC* promoter shown red. “Ex.” and “int.” are *α-SNAP_Rhg1_LC* exons and introns, respectively. LTR: long terminal repeat; GAG: group-specific antigen, RT: reverse-transcriptase. (E) Agarose gel showing 5’and 3’ *α-SNAP-RAC* junction products from the *rhg1-a* (low-copy, red dots) accessions: ‘Forrest’, PI 90763, PI 437654, PI 89772. No *α-SNAP-RAC* junctions detected from *rhg1-b* (high-copy; orange dots) accessions: PI 88788, PI 209332, or PI 548316. (F) Similar to E and using same template DNA samples, but PCR amplification of a WT (wild type) *α-SNAP* exon 1 – 2 distance, as in the Williams 82 reference genome. WT *α-SNAP* exon 1 to exon 2 distances outlined in *C* by an arrow and *.

An NCBI nucleotide BLAST of the unknown 4.77 kb region returned hits for conserved features of the Ty-1 copia retrotransposon superfamily. Notably, the multi-cistronic open reading frame (ORF) of this copia element was fully intact, and both 5’ and 3’ LTRs were present (Long Terminal Repeats; LTRs function as transcriptional promoters and terminators, respectively) (Havecker et al., 2004). Subsequently, we named this insert ‘*RAC*’ for *Rhg1* <u>a</u>lpha-SNAP <u>c</u>opia. Fig 1D shows the *rhg1-a α-SNAP*-*RAC* structure and Fig S2 provides the complete *RAC* nucleotide sequence and highlights the *α-SNAP_Rhg1_LC* intron 1 sequences directly flanking the *RAC-*integration. The *RAC* insertion effectively doubles the pre-spliced α-SNAP*_Rhg1_*LC mRNA transcript from 4.70 kb to 9.47 kb, yet *RAC* apparently splices out effectively, as all reported cDNAs of mature α-SNAP*_Rhg1_*LC transcripts do not contain any *RAC* sequences (Cook et al., 2014; Lee et al., 2015; Liu et al., 2017). The *RAC* ORF is uninterrupted and encodes a 1438 residue polyprotein with conserved copia retrotransposon motifs for GAG (group specific antigen) protease, integrase and reverse transcriptase (Peterson-Burch and Voytas, 2002; Havecker et al., 2004; Kanazawa et al., 2009). These conserved RAC polyprotein motifs are highlighted in Fig S3. Intriguingly, *RAC* integrated just 396 bp downstream of the *α-SNAP_Rhg1_LC* start codon, and in an anti-sense orientation (Fig 1D). The intact LTRs and uninterrupted ORF suggest that *RAC* integration could have been a relatively recent event, and that *RAC* may remain functional.

PI 89772 (used above) is one of seven *rhg1-a* and *rhg1-b* soybean accessions used to determine the HG type of SCN populations (Niblack et al., 2002). We subsequently tested for *RAC* insertions within the *Rhg1-*encoded *α-SNAP*(s) genes in the other six soybean accessions used in HG type tests. The *RAC* integration within the PI 89772-encoded *α-SNAP_Rhg1_LC* creates unique 5’ and 3’ sequence junctions within the *α-SNAP_Rhg1_LC* intron 1, and substantially increases the distance from *α-SNAP* exon 1 to exon 2 from ∼400 bp to ∼5,000 bp (Fig 1B and C). Therefore, to screen for *α-SNAP-RAC* insertions, we devised PCR assays specific for *RAC*-*α-SNAP* junctions, or for wild-type (uninterrupted) “WT junctions” separated by the genomic distances from exon 1 to exon 2 that are annotated in the soybean reference genome (Schmutz et al., 2010), as depicted in Fig 1C and 1D. Among all *rhg1-a* haplotype HG type test accessions (PI 90763, PI 89772, PI 437654 and PI 548402(Peking)-derived ‘Forrest’), both 5’ and 3’ *α-SNAP*-*RAC* junctions were detected (Fig 1D).

### *RAC* is absent from *rhg1-b* and single-copy *Rhg1_WT_* accessions

The *α-SNAP-RAC* junctions were absent from the *rhg1-b* accessions tested (PI 88788, PI 548316, PI 209332), which instead gave WT junction PCR products (Fig 1D). Because no SNPs exist at the WT junction primer sites across any of the *rhg1-b* repeats (Cook et al., 2014), absence of 5’ *RAC* junction and 3’ *RAC* junction PCR products for the *rhg1-b* accessions suggests that those accessions do not carry the *RAC* copia element in the first intron of their *α-SNAP* gene. Accession Williams 82 (Wm82, SCN-susceptible, *Rhg1* single-copy), the source of the soybean reference genome, also gave a product for the WT junction reaction, and no PCR product for a *RAC* integration within the *Rhg1_WT_* (single-copy) *α-SNAP* gene, consistent with the reference genome annotation (Schmutz et al., 2010) (Fig S1B). *RAC* absence from *rhg1-b* and WT *Rhg1* repeats is also consistent with previous studies that sub-cloned and Sanger-sequenced large-insert genomic fragments spanning entire *rhg1-b* and *Rhg1_WT_*-like repeats and noted no unusual insertions (Cook et al., 2012). Although all seven HG type test soybean lines were previously analyzed via WGS, the *RAC* insertion was evidently omitted from the four *rhg1-a* accession assemblies during Illumina short sequence read filtering that excludes repetitive genome elements, and hence from subsequent read mapping and assembly to the Williams 82 reference genome, which lacks the *RAC* insertion (Cook et al., 2014; Lee et al., 2015). The *RAC* insertion was apparently missed in other studies due to sequencing of post-splicing *Rhg1 α-SNAP* cDNAs (Bayless et al., 2016; Liu et al., 2017).

The WT junction PCR experiment also interrogated if *RAC* is present within each encoded *α-SNAP* gene of all three *rhg1-a* repeats. Among all *rhg1-a* accessions, no WT exon 1 to exon 2 junctions were detected, while uninterrupted WT junction product distances were present in all *rhg1-b* accessions and Wm82 as noted above (Fig 1E, F, Fig S1B). Hence *RAC* is apparently present within the *α-SNAP* of each *rhg1-a* repeat unit. To independently investigate the same question, previously available Illumina whole genome sequence data were queried (Cook et al., 2014). The read-depth for *RAC* was found to be 3-4 fold greater in *rhg1-a* accessions relative to the read-depth of the flanking DNA regions, or when compared to *RAC* read depth in *rhg1-b* accessions (Fig S4, SI Spreadsheet). Table 1 provides a summary of the *Rhg1* haplotype composition, copy number, resistance-type *α-SNAP* allele, and presence of normal vs. *RAC*-interrupted *α-SNAP* among the HG type test accessions and the Wm82 reference genome. Together, these findings indicate that the HG type test *rhg1-a* accessions contain the *RAC*-*α-SNAP* introgression, and that their *rhg1-a* repeats are ∼36.0 kb, as opposed to ∼31.2 kb for *rhg1-b* repeats and *Rhg1_WT_*. Although *RAC* is integrated in anti-sense orientation and close to *α-SNAP_Rhg1_LC* exon 1, *RAC* does not eliminate *rhg1-a* function, because the HG type test accessions are selected owing to their strong SCN-resistance, and, all have previously been shown to express α-SNAP*_Rhg1_*LC mRNA and protein (Cook et al., 2014; Bayless et al., 2016; Liu et al., 2017; Bayless et al., 2018). *RAC* presence within these *rhg1-a* lines may even be beneficial.

| Accession | *Rhg1*<br>Copies | SCN<br>Resistant | *Rhg1* WT<br>α-SNAP | *Rhg1* HC<br>α-SNAP | *Rhg1* LC<br>α-SNAP | WT<br>Junction | RAC<br>Junctions |
| --- | --- | --- | --- | --- | --- | --- | --- |
| Williams 82 | 1 | — | ✓ | — | — | ✓ | — |
| 'Forrest' | 3 | ✓ | — | — | ✓ | — | ✓ |
| PI 90763 | 3 | ✓ | — | — | ✓ | — | ✓ |
| PI 437654 | 3 | ✓ | — | — | ✓ | — | ✓ |
| PI 89772 | 3 | ✓ | — | — | ✓ | — | ✓ |
| PI 548316 | 7 | ✓ | ✓ | ✓ | — | ✓ | — |
| PI 88788 | 9 | ✓ | ✓ | ✓ | — | ✓ | — |
| PI 209332 | 10 | ✓ | ✓ | ✓ | — | ✓ | — |

### *RAC* is not identical to other copia elements but the *RAC-*like copia subfamily is common in soybean

Copia retrotransposons frequently attain high copy numbers in plant and animal genomes, therefore, we assessed the abundance of *RAC* and *RAC*-like copia elements in the soybean genome (Du et al., 2010; Du et al., 2010; Zhao and Ma, 2013). SoyTE, the soybean transposon database, has recorded over 32,000 transposable elements, including nearly 5,000 intact retrotransposons (Du et al., 2010). We queried the SoyTE database via Soybase.org using a *RAC* nucleotide sequence BLASTN search, but no intact or high identity hits were returned. However, a similar BLASTN search against the Wm 82 soybean reference genome at Phytozome.org (Goodstein et al., 2012) returned 146 sequences. These 146 hits spanned all 20 *G. max* chromosomes and included several intact elements of high nucleotide identity with *RAC*, as well as numerous short length matches which likely represent fragments of inactive elements (Table S1). We then constructed a nucleotide-based phylogenetic tree of the soybean *RAC* family using just one *RAC-*family element from each soybean chromosome (Chr) (Fig 2A). This analysis used the element from each chromosome that was most similar to *RAC*, as well as a previously reported copia retrotransposon (TGMR) residing near the soybean *Rps1-k* resistance gene (Bhattacharyya et al., 1997) and the highest *RAC-*identity element match from the common bean (*Phaseolus vulgaris*) genome (Fig 2A). The two *RAC-*like elements, from Chr10 and Chr18, had 99.7% and 97.6% nucleotide identity with *RAC*, respectively (Fig 2A, Table S1). Moreover, the Chr10 element retained an intact ORF and both LTRs. The near-perfect nucleotide identity with *RAC* (99.7%; 4464/4477 positions) suggests the Chr10 element as a possible source for the retrotransposition event that created the *RAC* introgression within *α-SNAP_Rhg1_LC*. The above-noted WGS read-depth analysis of soybean accessions (Fig S4, SI Spreadsheet) also found that 3 of 10 examined *rhg1-b* accessions gave a read-depth of zero or close to zero for *RAC* (with one mismatch allowed), indicating absence of the *RAC* or a close homolog at the Chr 10 locus in some soybean accessions. Like *RAC*, the Chr18 element was also integrated anti-sense within a host gene *Glyma.18G268000* (a putative leucine-rich repeat receptor kinase). We further noted that the Chr20 *RAC*-like element (82% identical) was intronically positioned within *Glyma.20G250200* (BAR-domain containing protein) (Table S1). That multiple intact and highly similar *RAC*-like elements are in soybean suggests that this retrotransposon family was recently active.

**Fig 2.**
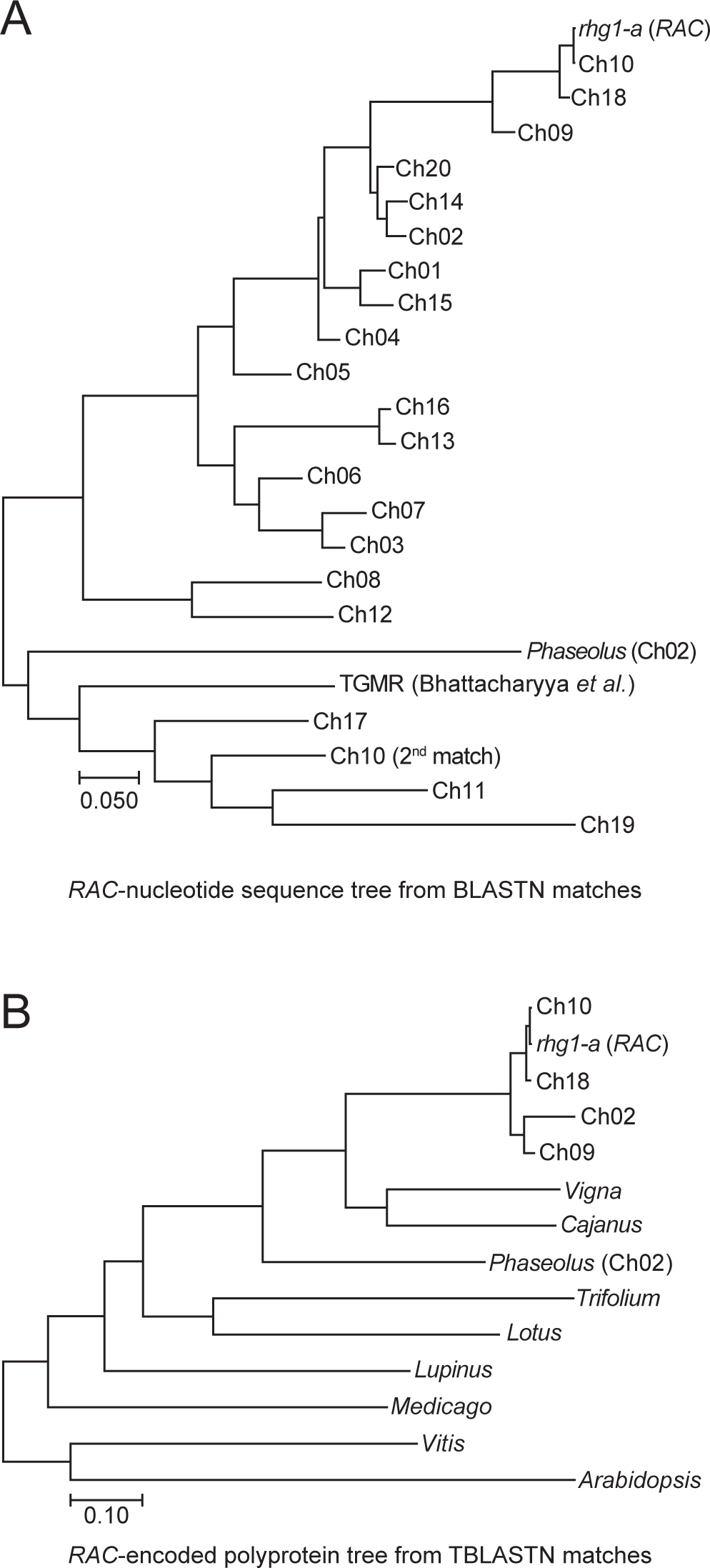
The *RAC*-like subfamily of copia retrotransposons is common in soybean and other legumes. (A) Maximum likelihood phylogenetic tree of *RAC*-like element nucleotide sequences from soybean. The top hit from each soybean chromosome was included, as was the known soybean retrotransposon “TGMR” and the top *RAC*-like match from *Phaseolus* (common bean). (B) Similar to *A*; a maximum-likelihood tree, but using the *RAC*-encoded polyprotein sequences from the four most similar soybean *RAC*-like elements, and the most similar element matches from the indicated plant species.

Subsequent work examined if *RAC-*family elements are present among other plant species. We performed a TBLASTN search of the *RAC*-encoded polyprotein at NCBI and obtained numerous hits against multiple species, including *Arabidopsis*, *Cajanus cajans* (pigeon pea), *Vigna angularis* (adzuki bean), *Phaseolus vulgaris* (common bean)*, Lupinus angustifolius* (blue lupin), *Medicago truncatula*, and clover (*Trifolium subterraneum*). Similar to Fig 2A, a phylogenetic tree using MEGA (muscle alignment) and the *RAC-*family polyprotein sequences of these different plant species was constructed (Kumar et al., 2016) (Fig 2B). Together, these findings demonstrate that the *RAC-*like copia members are not only common in *Glycine max*, but also in other legumes.

### *RAC* is present within only a subclass of the soybean accessions that have the previous SoySNP50K-predicted *rhg1-a* signature

Although we detected *RAC* in all four *rhg1-a* accessions that are used for HG type determination, we sought to determine if the *RAC* insertion is universal among all *rhg1-a-* containing accessions. Recently, the USDA soybean collection (∼20,000 accessions) was genotyped using a 50,000 SNP DNA microarray chip – the SoySNP50K iSelect BeadChip (Song et al., 2015). We searched for and found a SNP on the SoySNP50K chip that detects *RAC*. Using the SoySNP50K browser at Soybase.org (Soybase.org/snps/), we found a SNP (ss715606985, G to A) that in the Wm82 soybean reference genome was assigned to the Chr10 *RAC-*family element (99.7% nucleotide identity with *RAC*). However, we noted that this ss715606985 SNP is a perfect match to the sequence of *RAC* within *α-SNAP_Rhg1_LC* (Fig S5A). Using this SNP marker for *RAC*, we then calculated the ss715606985 SNP prevalence among all USDA accessions, and found that the SNP is rare – only 390 of 19,645 accessions (∼2.0%) were putative *RAC^+^* lines homozygous for the SNP (Fig 3A). The *RAC-*SNP (ss715606985) was then directly tested as a marker for the *α-SNAP*-*RAC* event using the PCR assays described in Fig 1, which test for *α-SNAP*-*RAC* junctions and normal *α-SNAP* exon 1-2 distances. We randomly selected several accessions with SNP-signatures of *rhg1-a* that were positive or negative for the *RAC*-SNP, and found that SNP presence correlated perfectly with *RAC*-*α-SNAP* junction detection, while accessions lacking the *RAC*-SNP had normal *α-SNAP* exon 1-2 distances indicative of no inserted DNA (Fig 3B).

**Fig 3.**
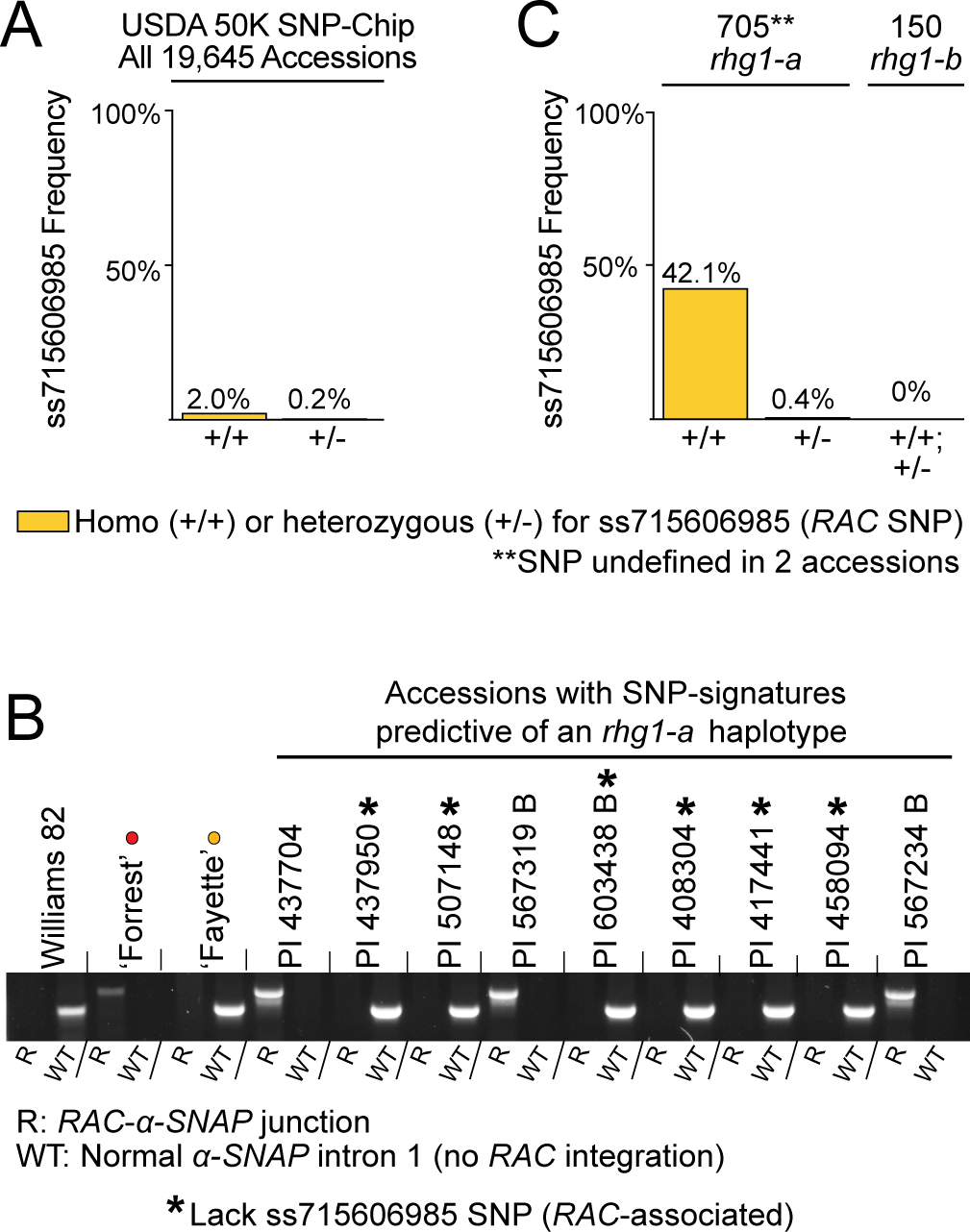
*RAC* is present within a subclass of *rhg1-a*-signature soybean accessions. (A) Frequency of *RAC*-associated SNP, ss715606985, among 19,645 SoySNP50K-genotyped USDA soybean accessions. (B) Agarose gel showing PCR detection of *α-SNAP-RAC* junctions or WT *α-SNAP* exon 1-2 distances among *rhg1-a*-signature accessions positive or negative for the *RAC*-SNP, ss71560698. Williams 82 (*Rhg1_WT_*), ‘Forrest’ (*rhg1-a*) and ‘Fayette’ (*rhg1-b*) included as controls; *Rhg1* haplotypes color coded with dots as in Fig 1. An * denotes an *rhg1-a*-signature accession lacking the *RAC*-SNP. (C) Frequency of *RAC*-associated SNP among all USDA Glycine *max* accessions with consensus SNP signatures of *rhg1-a* or *rhg1-b* haplotypes.

We next examined *RAC* presence among all *Glycine max* USDA accessions with the SoySNP50K SNP-signatures of *rhg1-a* or *rhg1-b* haplotypes, as reported by Lee *et al*.(Lee et al., 2015). The multi-SNP SoySNP50K signatures for *rhg1-a* and *rhg1-b* (Lee et al., 2015) are present in 705 and 150 *Glycine max* accessions respectively, out of 19,645 USDA accessions; these SoySNP50K signatures are provided in Fig S5B. We found that 42% (299 of 705) of accessions with *rhg1-a* SNP-signatures and 0% (0 of 150) of accessions with *rhg1-b* SNP-signatures carry the *RAC* ss715606985 SNP (Fig 3C). That the *RAC-*SNP was absent from all *rhg1-b* signature accessions is consistent with the PCR screens of Fig 1E and F, which indicated that no *rhg1-b* HG type test accession contained *RAC*-*α-SNAP* junctions. A flow-chart is available as Supplementary Fig S6 that summarizes the above findings and additional work presented below.

Because only 299 of the 390 accessions with the *RAC*-SNP had a perfect-match *rhg1-a* SoySNP50K SNP signature (Fig 3C), we investigated the *Rhg1* SNP signature of the remainder. To avoid false positives, the *rhg1-a* SNP signature of (Lee et al., 2015) uses 14 SNPs that extend to 11 kb and 54 kb beyond the edges of the ∼30 kb *Rhg1* repeat. Relaxing the stringency that required 14 perfect matches, we found that 83 of the remaining 91 *RAC* ^+^ accessions carry a perfect match with the four *rhg1-a* SNP markers that map within the *Rhg1* repeat or within <5 kb of the edge of the *Rhg1* repeat. 86 of 91 have only a single reliably called SNP that varies from the *rhg1-a* consensus (SI Spreadsheet). Together, the combined findings indicate that the *α-SNAP-RAC* integration is only present within *rhg1-a* haplotypes, and that *RAC* retrotransposition may have occurred within a subset of the *Rhg1^+^* population after *Rhg1* divergence into the distinctive high- and low-copy haplotype classes.

### The *RAC-*SNP allows more accurate prediction of *rhg1-a* presence

The above finding that a few hundred of the 705 USDA accessions with the previously identified *rhg1-a* SNP signature apparently do not contain *RAC* was surprising, given that all four of the *rhg1-a* HG type test accessions do contain *RAC*-*α-SNAP* junctions. However, it was possible that these non-*RAC* containing accessions, despite a consensus SNP-signature predicting an *rhg1-a-*haplotype, might not truly carry an *rhg1-a* resistance haplotype. *rhg1-a* and *rhg1-b* repeats encode distinct *Rhg1 α-SNAP* alleles, thus, we cloned and sequenced the genomic *Rhg1 α-SNAP* regions from several non-*RAC rhg1-a* SNP-signature accessions and detected coding sequences for either *rhg1-b* (*α-SNAP_Rhg1_HC*) or *Rhg1_WT_* (*α-SNAP_Rhg1_WT*) alleles (Fig S5C). None of these accessions encoded *α-SNAP_Rhg1_LC* and thus, they were not *rhg1-a* (Fig S5C). These findings indicate that the consensus *rhg1-a* SNP-signature, while useful, is not a perfect predictor of accessions carrying actual *rhg1-a* resistance haplotypes. Rather, combined presence of the ss715606985 SNP for *RAC* and a near-consensus *rhg1-a* signature is a more accurate predictor of accessions that truly carry *rhg1-a* resistance. Additionally, these data suggest that accessions carrying *rhg1-b* resistance haplotypes can share the SNP signatures of *rhg1-a* accessions. We again refer readers to the flow-chart (Supplementary Fig S6) that summarizes these and other findings.

### *RAC* presence correlates with a stronger SCN-resistance profile and the co-presence of other loci that augment *rhg1-a* resistance

*rhg1-a* (*Rhg1* low-copy) loci encode unique *α-SNAP_Rhg1_LC* alleles, however, robust *rhg1-a* resistance requires the co-presence of *Rhg4*, and the *α-SNAP Ch11-IR* allele bolsters *rhg1-a* resistance further (Liu et al., 2012; Lakhssassi et al., 2017; Bayless et al., 2018; Patil et al., 2019). We sought to compare the SCN resistance profiles of the *rhg1-a* signature accessions with *RAC* to those without *RAC* (which are not true *rhg1-a*), to assess how the two groups match what is known about *rhg1-a* resistance. Previously, Arelli, Young, and others obtained SCN resistance phenotype data, across multiple trials and with various SCN populations, for at least 573 different USDA accessions that are now known to carry the SoySNP50K signatures suggestive of *rhg1-a* (Anand, 1984; Hussey et al., 1991; Young, 1995; Diers et al., 1997; Arelli et al., 2000; Lee et al., 2015). We used these available SCN resistance data from the USDA GRIN database to compare the resistance profiles of *rhg1-a*-signature accessions which did or did not have the ss715606985 (*RAC*) SNP signature. Any accession that scored as “R” (resistant) in any single SCN trial was placed into the broad category “R”. Likewise, any accession that scored “MR” (moderately resistant) or “MS” (moderately susceptible) in any trial, with no higher resistance scores in other trials, was placed into those respective categories. Only accessions that scored susceptible (“S”) across all trials were placed into the “S” category. Consistent with previous reports that *rhg1-a* accessions possess broad and robust resistance(Concibido et al., 2004; Vuong et al., 2015; Kadam et al., 2016), 91% (51/56) of the accessions in the “R” group were positive for the ss715606985 ^+^ *RAC* SNP (Fig 4A). The frequency of *RAC* presence was substantially lower among the more susceptible phenotypic classes (Fig 4A) while the majority of the ss715606985 ^−^ (no *RAC*) accessions scored either “S” or “MS”. As was noted above, none of the non-*RAC* accessions that we examined had *rhg1-a* type resistance (Fig 3B, 3C, Fig S5C). The phenotype scores and relevant SNP markers for all 573 of these SCN-phenotyped *rhg1-a* SNP signature accessions are provided as a spreadsheet in the SI Data.

**Fig 4.**
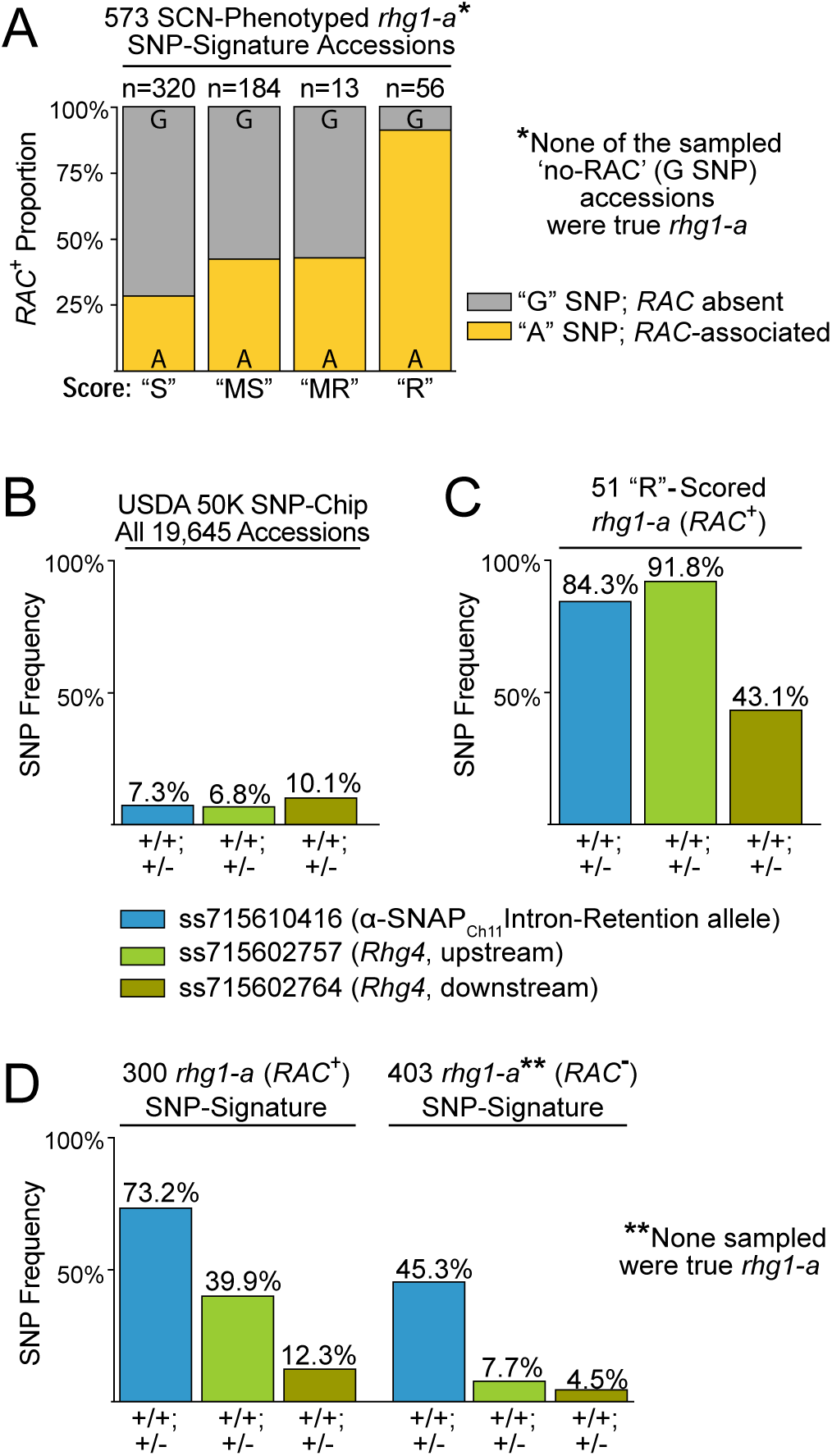
*RAC* presence correlates with a stronger SCN-resistance profile and the co-presence of other loci that augment *rhg1-a* resistance. (A) Proportion of *RAC* ^+^ (ss715606985 A SNP) vs. *RAC* ^−^ (G SNP) accessions among 573 SCN-phenotyped soybeans with consensus SoySNP50K SNP signatures predictive of *rhg1-a*. *Note that none of the sampled *RAC* ^−^ (G SNP) accessions had *rhg1-a* (none encoded *α-SNAP_Rhg1_LC*). “S”: susceptible in all trials, “MS”: moderately susceptible in at least one trial, “MR”: moderately resistant in at least one trial, “R”: resistant in at least one trial. Fisher’s Exact Test pairwise comparisons: “R-MR” (p = 2.6E-4), “R-MS” (p = 2.3E-11), “R-S” (p = 2.2E-16), “MR-MS” (p = 1.0), “MR-S” (p = 0.25), “MS-S” (p = 2.4E-3). (B) Frequency of SNPs associated with *Rhg4* (ss715602757, ss715602764) or the Chromosome 11-encoded α-SNAP-Intron Retention (α-SNAP_Ch11_IR) allele, ss71559743 among 19,645 USDA accessions. (C) Frequency of the *Rhg4* and *α-SNAP_Ch11_IR* associated SNPs among the 51 “Resistant” scored *RAC* ^+^ *rhg1-a*-signature accessions. (D) Frequency of the *Rhg4* and *α-SNAP_Ch11_-IR* associated SNPs among all *RAC* ^+^ (300) or *RAC*^−^ (403) USDA *G. max* accessions with consensus SNP signatures predictive of *rhg1-a* (705 total; 2 accessions undefined for ss715606985 SNP).

In the above analysis (Fig 4A), some of the *RAC^+^* (ss715606985^+^) accessions, which are *rhg1-a*, had scored as “S” or “MS”. This seemed likely to be because they lack a resistance-conferring allele at *Rhg4* and/or the resistance-enhancing allele of the Chr 11-encoded *α-SNAP* (*α-SNAP_Ch11_-IR*) (Liu et al., 2012; Lakhssassi et al., 2017; Bayless et al., 2018). Accordingly, we investigated if the SCN resistance phenotype scores also correlated with co-presence of those loci. None of the SoySNP50K markers resides within the *Rhg4* gene but we noted that the two SoySNP50K SNPs that most closely flank the *Rhg4* locus are rare among USDA accessions (Fig 4B; ss715602757, ss715602764), and one or both of these SNPs is present in the *Rhg4*-containing HG type test lines. We also used a SNP, ss715610416, previously associated with the Chr11 *α-SNAP* intron-retention allele (*α-SNAP_Ch11_-IR*) (Bayless et al., 2018). Among the 51 *RAC*^+^ accessions with an SCN resistance score of “R”, we found that SNPs associated with both *α-SNAP_Ch11_-IR* and *Rhg4* were enriched ∼10-fold, as compared to the entire USDA collection (Fig 4 B,C). Additionally, among the 705 USDA accessions with SoySNP50K signatures predictive of *rhg1-a*, we found that the *Rhg4* and *α-SNAP_Ch11_-IR* SNPs were enriched among the *RAC*^+^ accessions relative to the *RAC*^−^ accessions (Fig 4D). Thus, the heightened SCN resistance of the *RAC*-positive (ss715606985 ^+^) *rhg1-a*-signature accessions is consistent with previous reports, and as expected, the resistance is associated with the co-presence of additional loci like *Rhg4* and *α-SNAP_Ch11_-IR* (Vuong et al., 2015; Kadam et al., 2016; Lakhssassi et al., 2017; Bayless et al., 2018; Patil et al., 2019).

### The *rhg1-a RAC* element has intrinsic transcriptional activity

While the *RAC*-SNP apparently identifies true *rhg1-a* resistance sources, possible impacts of the *RAC* element itself on *α-SNAP_Rhg1_LC* expression remained to be explored. Typically, eukaryotic cells silence transposable elements using small RNA-directed DNA methylation pathways, and this can also silence adjacent genes (McCue et al., 2012; Kim and Zilberman, 2014). Since the α-SNAP*_Rhg1_*LC mRNA transcript and protein are readily detected in *RAC-*containing soybeans, *RAC* does not, at least constitutively, eliminate *α-SNAP_Rhg1_LC* expression (Cook et al., 2014; Bayless et al., 2016; Liu et al., 2017; Bayless et al., 2018). Nonetheless, we examined DNA methylation at the *rhg1-a α-SNAP-RAC* junction, as well as transcriptional activity of the *rhg1-a RAC* element. The restriction enzyme McrBC cleaves only methylated DNA, so potentially methylated DNA regions may be assessed via McrBC digestion and subsequent attempted PCR across areas of interest. After genomic DNAs from ‘Forrest’ and ‘Peking’ (PI 548402) were treated with McrBC, both the 5’ and 3’ borders of the *α-SNAP*-*RAC* did not PCR amplify relative to the mock treated controls, indicating the presence of methylated cytosines at the *α-SNAP*-*RAC* junctions (Fig 5A).

**Fig 5.**
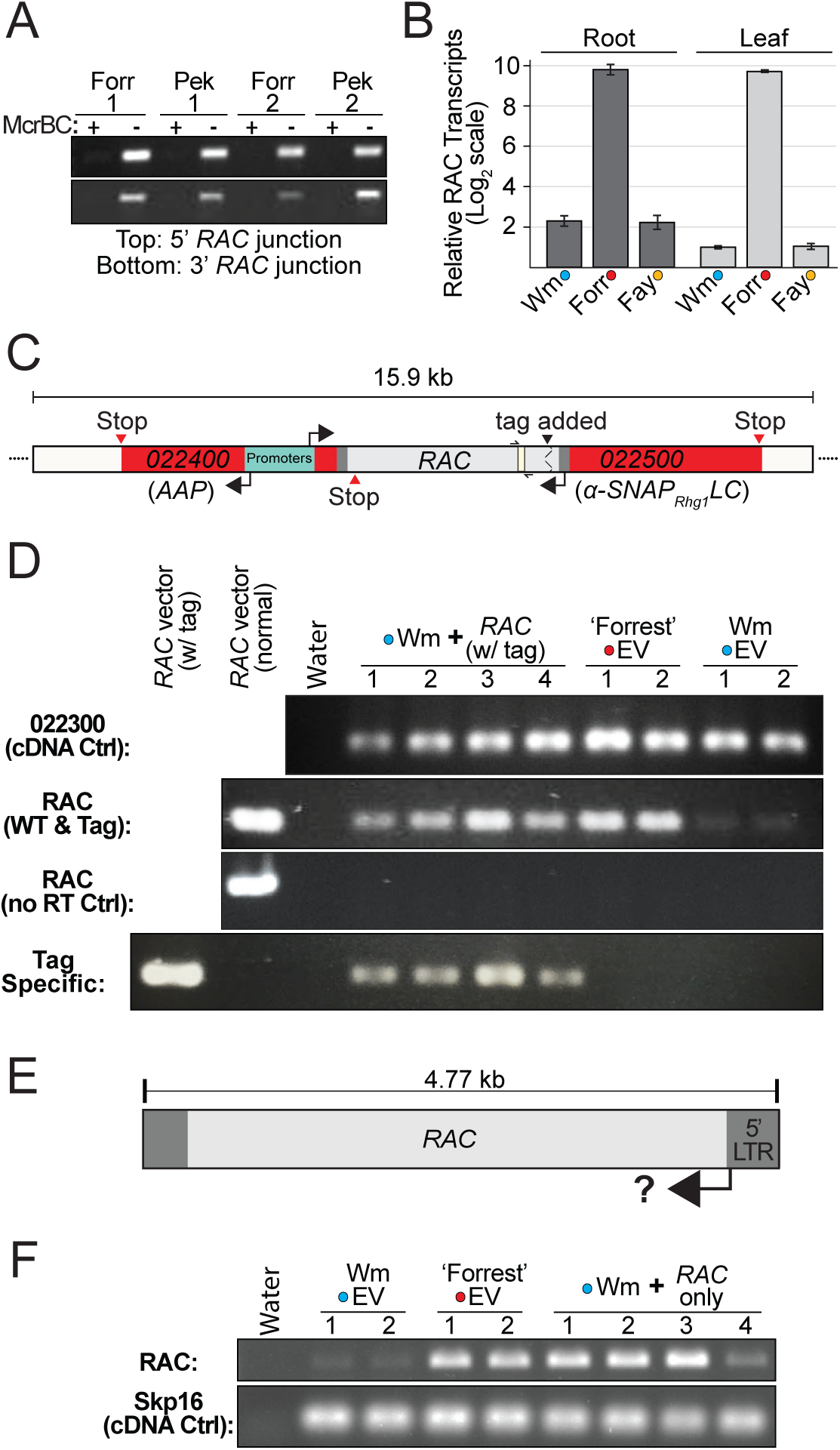
The *rhg1-a RAC* element is methylated but has intrinsic transcriptional activity. (A) Agarose gel showing PCR amplicons for *α-SNAP*-*RAC* regions from McrBC treated (+) or mock treated (-) genomic DNAs from ‘Forrest’ (Forr) or ‘Peking’ (Pek, PI 548402) roots. (B) qPCR analysis of mRNA transcript abundance for *RAC* and similar *RAC*-like elements, in leaf or root tissues of Williams 82 (Wm; *Rhg1_WT_*), ‘Forrest’ (Forr; *rhg1-a*) or ‘Fayette’ (Fay; *rhg1-b*). Colored dots indicate *Rhg1* haplotype as in Fig 1. Normalized *RAC* transcript abundances are presented relative to the mean abundance of *RAC* transcript for Williams 82 leaf samples. Y-axis uses log_2_ scale. (C) Schematic showing unique nucleotide tag addition to an otherwise native *α-SNAP*-*RAC* cassette. This construct contains native flanking *Rhg1* sequence including *Glyma.18G022400* (transcribes from the bidirectional *α-SNAP* promoter) and 1.8 kb upstream, as well as 4.7 kb of downstream *RAC* flanking sequence (∼1.0 kb after the *α-SNAP_Rhg1_LC* termination codon). The *RAC* region detected and amplified via qPCR or RT-PCR is colored ivory and flanked by half-arrows. (D) Agarose gel of RT-PCR cDNAs of ‘Forrest’ or Wm 82 transgenic roots transformed with an empty vector (EV) or the native tagged *α-SNAP*-*RAC* construct. Tag primers amplify only the modified *α-SNAP*-*RAC* while the normal *RAC* primer set amplifies both endogenous *RAC*-like transcripts as well as the tagged *α-SNAP*-*RAC* transgene. *Glyma.18G022300* mRNA transcript used as a cDNA quality and loading control; no RT (reverse transcriptase) ctrl verifies absence of amplifiable genomic DNA. (E) Schematic showing the subcloned 4.77 kb *RAC* expression cassette tested in F. (F) Like D, but ‘Forrest’ or Wm 82 roots transformed with empty vector or the 4.77 kb *RAC* element (all flanking *Rhg1* sequence context removed).

Because *RAC* has both LTRs and an intact ORF, we tested for transcription of *RAC* in the *rhg1-a* soybean genotype ‘Forrest’ as compared to ‘Fayette’ (*rhg1-b*) and Wm82 (*Rhg1_WT_*). Fayette and Wm82 do not carry the *Rhg1 α-SNAP-RAC* but do carry other *RAC*-like copia elements that match the qPCR primers used (diagrammed in Fig 5C). qPCR analysis of cDNAs from root or leaf tissues indicated that mRNA transcripts from *RAC* or *RAC-*like sequences were ∼200-fold higher in ‘Forrest’ than in Wm82 or ‘Fayette’ (*rhg1-b*) (Fig 5B). This suggested but did not firmly demonstrate that the *Rhg1*-embedded *RAC* is the primary source of the detected transcript, because *RAC* has high nucleotide identity with other *RAC-*like elements (Fig 2A) whose activity may also vary between accessions.

We conducted additional tests for transcription of *α-SNAP*-*RAC* by transforming Wm82 roots with a ∼15 kb cloned segment of native *α-SNAP*-*RAC* genomic DNA (including the upstream *Glyma.18G024400 Rhg1* gene, which shares the same bidirectional promoter; depicted in Fig 5C). Importantly, we engineered this otherwise native *α-SNAP*-*RAC* cassette with a unique nucleotide tag to distinguish between transgene derived transcripts vs. transcripts from other *RAC-*family elements in the genome (Fig 5C). Low abundance of *RAC* transcripts in Wm82 roots relative to ‘Forrest’ roots had been documented (Fig 5B), so RT-PCR of Wm82 readily visualized *RAC*-specific transcript expression from the transgenically introduced construct. In control roots, sharp contrasts in *RAC* expression were again observed between ‘Forrest’ roots and Wm82 roots transformed with empty vector (Fig 5D). But Wm82 roots transformed with the uniquely tagged *α-SNAP*-*RAC* transgene had substantially elevated *RAC* transcripts compared to isogenic Wm82 controls, as indicated by a primer pair that amplifies all *RAC* sequences (native or tagged), and by a primer pair that amplifies only the uniquely tagged *α-SNAP*-*RAC* transcript (Fig 5D). Controls using template samples prepared without reverse transcriptase verified successful DNAase treatment of cDNA preparations (Fig 5D). We further tested the activity of the *RAC* promoter itself by constructing a native 4.77 kb *RAC* element cassette divorced from the flanking *Rhg1* DNA (Fig 5E), which we then transformed into Wm82. Similar to Fig 5D, the native 4.77 kb *RAC* transgene substantially increased *RAC* transcript abundance in Wm82 roots, relative to empty vector controls (Fig 5F). Together, these findings demonstrate that presence of the *rhg1-a* locus *RAC* can substantially elevate *RAC* mRNA transcripts, and that *RAC* itself possesses intrinsic promoter activity. The findings suggest that the high *RAC* transcript abundance observed in ‘Forrest’ (*rhg1-a*), but not ‘Fayette’ (*rhg1-b*) or Wm 82 (single-copy *Rhg1*), is likely to be derived from the *rhg1-a* locus *RAC* insertion. These findings also support the possibility that *RAC* may retain the potential to promote transposition.

### α-SNAP*_Rhg1_*LC protein is expressed despite *RAC* presence

TEs can influence the expression of host genes. Because *RAC* is present in *rhg1-a* accessions previously chosen for use in agricultural breeding due to their strong SCN-resistance, *RAC* presence may benefit *rhg1-a*-containing soybeans. In light of *RAC’s* anti-sense orientation and close proximity to the *Rhg1 α-SNAP_Rhg1_LC* promoter, we sought to examine if *RAC* influences α-SNAP*_Rhg1_*LC protein expression. We were not able to compare expression of α-SNAP*_Rhg1_*LC between native *rhg1-a* loci that do or do not contain *RAC*, because no *rhg1-a* accessions without *RAC* have been identified and deleting *RAC* from all *Rhg1* repeats of a *rhg1-a* accession would not be trivial. Therefore, we left intact or removed the 4.77 kb *RAC* insertion from the native *α-SNAP*-*RAC* construct used for Fig 5 (Fig 6A) and then examined α-SNAP*_Rhg1_*LC protein abundance in transgenic Wm82 roots carrying the respective constructs. Immunoblotting was conducted using previously described α-SNAP*_Rhg1_*LC-specific and WT α-SNAP-specific antibodies (Bayless et al., 2016). The results of a representative experiment are shown in Fig 6B. Across multiple experiments containing independently transformed roots, the constitutive expression of α-SNAP*_Rhg1_*LC protein was highly variable, regardless of *RAC* presence/absence. However, with respect to constitutive expression of α-SNAP*_Rhg1_*LC protein we observed no requirement for *RAC* nor any obvious detrimental impact of *RAC* (Fig 6B).

**Fig 6.**
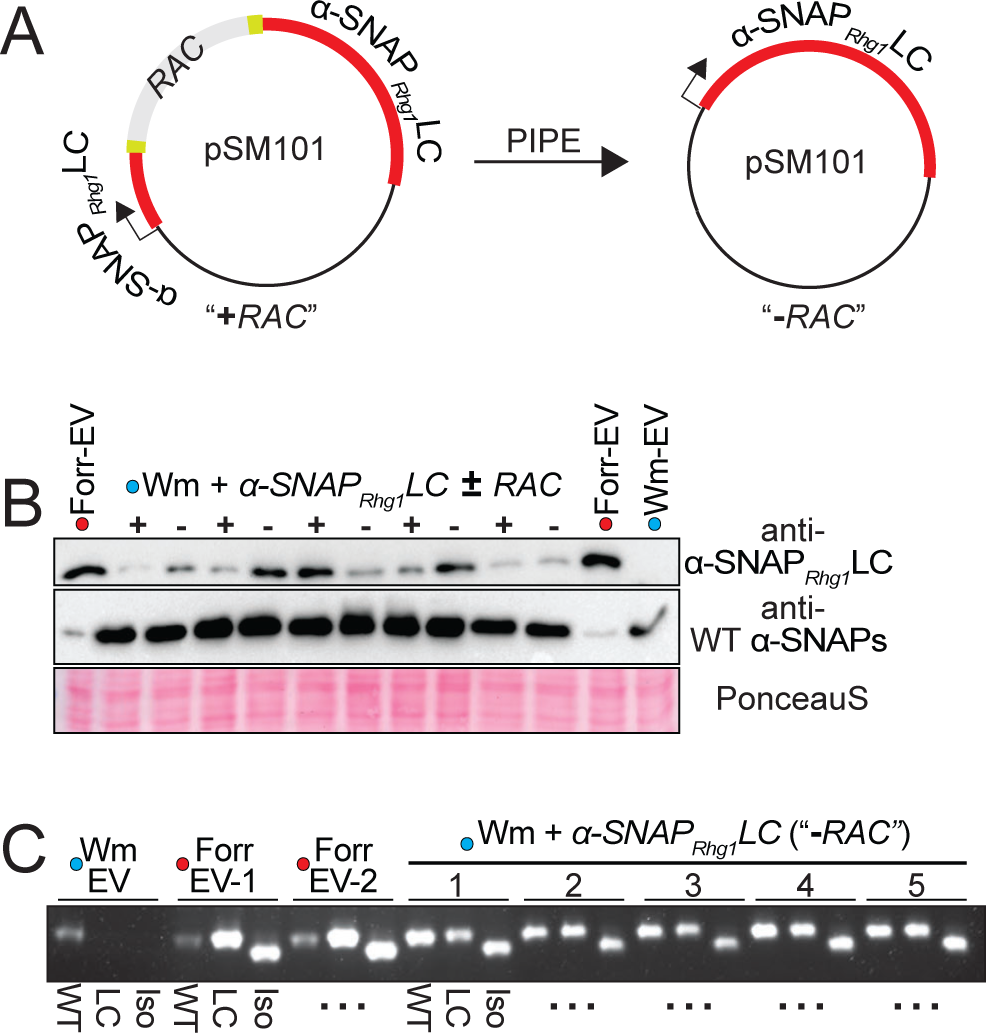
α-SNAP*_Rhg1_*LC protein is expressed despite *RAC* presence. (A) Schematic showing PIPE-mediated removal of *RAC* from the native *α-SNAP*-*RAC* construct, pSM101. (B) Immunoblots of independent ‘Forrest’ or Wm82 transgenic root lysates using previously described antibodies for α-SNAP*_Rhg1_*LC or WT α-SNAP proteins. “+” denotes *α-SNAP*-*RAC* transformation, “-” indicates *α-SNAP_Rhg1_LC* (*RAC* removed) transformed, and EV is empty vector transformed. Ponceau S staining serves as a loading control. (C) Agarose gel showing RT-PCR amplification of mature α-SNAP*_Rhg1_*LC transcript isoforms from roots of Wm 82 or ‘Forrest’ transformed with *α-SNAP*-*RAC* (+), or a native *α-SNAP_Rhg1_LC* cassette with *RAC* removed (**-**), or an empty vector control. WT refers to primers specific for WT α-SNAP transcripts, LC detects full length α-SNAP*_Rhg1_*LC transcripts, while “Iso” amplifies a previously described α-SNAP*_Rhg1_*LC alternative transcript isoform that splices out 36 bp (Cook et al., 2014).

Previously, we reported that native α-SNAP*_Rhg1_*LC mRNA transcripts include an alternative splice product (Cook et al., 2014; Bayless et al., 2016). Because TEs can influence host mRNA splicing (Krom et al., 2008), we also examined how *RAC* influenced splicing of the known α-SNAP*_Rhg1_*LC alternative transcript. As above, we generated transgenic roots of Wm82 containing either a *RAC* ^+^ *α-SNAP_Rhg1_LC* native genomic segment or a version with *RAC* precisely deleted (Fig 6A). We then generated cDNAs and performed RT-PCR with primer sets specific for either the full length or shorter splice isoform. As shown by agarose gel electrophoresis, *RAC* presence was not required for alternate splicing of this α-SNAP*_Rhg1_*LC isoform (Fig 6B).

### α-SNAP*_Rhg1_*LC hyperaccumulates at SCN infection sites

We also examined infection-associated α-SNAP*_Rhg1_*LC protein expression in non-transgenic soybean roots that carry the native *rhg1-a* locus. We previously reported that during *rhg1-b*-mediated SCN-resistance, α-SNAP*_Rhg1_*HC abundance is elevated ∼12-fold within syncytial cells (SCN feeding sites) relative to adjacent non-syncytial cells (Bayless et al., 2016). To test whether the *RAC*-containing *rhg1-a* follows a similar expression pattern during the resistance response, the present study examined α-SNAP*_Rhg1_*LC abundance at SCN infection sites in soybean variety ‘Forrest’ using SDS-PAGE and immunoblots. Using the aforementioned α-SNAP*_Rhg1_*LC-specific antibody, we detected increased α-SNAP*_Rhg1_*LC accumulation within tissues enriched for SCN feeding sites, while expression was barely detectable in mock-treated roots (Fig 7A). As previously reported, NSF proteins were also increased in SCN-infested roots, albeit less prominently (Fig 7A) (Bayless et al., 2016). Thus, even in *RAC* presence, *rhg1-a* haplotypes drive an expression pattern of α-SNAP*_Rhg1_*LC similar to that observed for *rhg1-b* and α-SNAP*_Rhg1_*HC.

**Fig 7.**
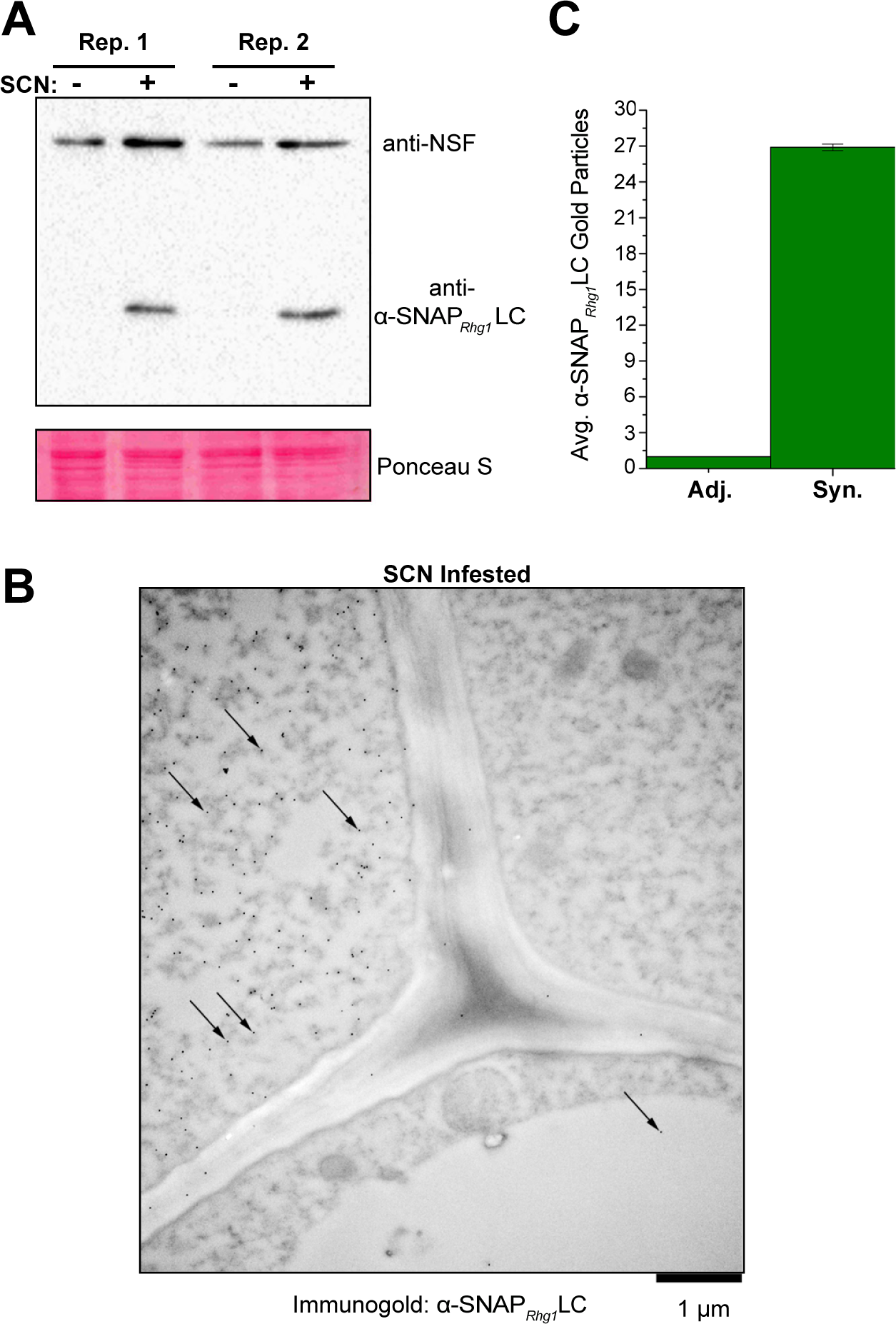
α-SNAP*_Rhg1_*LC hyperaccumulates at SCN infection sites in low-copy *rhg1-a* soybean accession ‘Forrest’. (A) Immunoblot of non-transgenic ‘Forrest’ root samples from SCN-infested root regions (SCN +) harvested 4 d after SCN infection, or similar regions from mock-inoculated controls (SCN -). Blot was probed simultaneously with anti-α-SNAP*_Rhg1_*LC and anti-NSF polyclonal antibodies. Ponceau S staining before blotting served as a loading control. (B) Representative electron microscope image (7 dpi) showing anti-α-SNAP*_Rhg1_*LC immunogold signal in SCN-associated syncytium cells from ‘Forrest’ roots. Arrows highlight only some of the 15 nm immunogold particle dots. Frequent α-SNAP*_Rhg1_*LC signal was observed in syncytium cells (upper left, “Syn”) but rare in adjacent cells (upper right and bottom, “Adj.”). CW, cell wall; M, mitochondrion; Vac, vacuole. Bar = 1 μm. (C) Mean and SEM of α-SNAP*_Rhg1_*LC gold particle abundance in syncytia, normalized to the count from adjacent cells in the same image. Anti-α-SNAP*_Rhg1_*LC immunogold particles were counted for one 9 μm^2^ area within cells having syncytium morphology and in a region with the highest observable signal in directly adjacent cells with normal root cell morphology (large central vacuole). Data are for 23 images (11 and 12 root sections respectively, from two experiments), for root sections 7 days after inoculation.

To more precisely locate the α-SNAP*_Rhg1_*LC increases, SCN-infested root sections were imaged using transmission electron microscopy and immunogold labelling of bound α-SNAP*_Rhg1_*LC-specific antibody. Syncytium-specific accumulation of α-SNAP*_Rhg1_*LC protein was observed and quantified in root sections taken 7 days after SCN inoculation (Fig 7B, C). The average increase of immunogold particles per equal area of adjacent non-syncytial root cells (cells still carrying a large central vacuole) was ∼25-fold (Fig 7C). In control experiments, EM sections from mock-inoculated roots (no SCN) exhibited no immunogold signal above background (Fig S7A). Similarly, no immunogold signal above background was observed when secondary antibody and all other reagents were used but the primary antibody was omitted (Fig S7B). The specificity of the antibody for α-SNAP*_Rhg1_*LC protein was previously demonstrated (signal for recombinant α-SNAP*_Rhg1_*LC protein or total protein from roots with *rhg1-a*, no signal for α-SNAP*_Rhg1_*HC protein or total protein from roots with *rhg1-b* or *Rhg1_WT_*) [30]. The above results, similar to the previously observed ∼12-fold increase reported for the α-SNAP*_Rhg1_*HC in syncytia from *rhg1-b* roots, indicate that α-SNAP*_Rhg1_*LC protein abundance is also elevated within syncytia upon SCN infection [30]. Collectively, these findings demonstrate that while a potentially active retrotransposon (*RAC*) has integrated within the important *rhg1-a α-SNAP_Rhg1_LC* resistance gene, and its presence correlates with *rhg1-a* haplotypes preferred for SCN resistance breeding, no negative impacts of *RAC* on α-SNAP*_Rhg1_*LC mRNA or protein expression were detected.

## Discussion

*Rhg1* is the principal SCN resistance locus in commercially grown soybeans. The increasing occurrence of SCN populations that at least partially overcome the overwhelmingly utilized “PI 88788-type” *rhg1-b* resistance source is an important concern for soybean breeders and growers (McCarville et al., 2017), (www.thescncoalition.com). Alternating use of different *Rhg1* haplotypes should help bolster and preserve resistance against these virulent SCN populations (Brucker et al., 2005; Niblack et al., 2008)(www.thescncoalition.com). In this study we report that the other *Rhg1* haplotype available for SCN control, *rhg1-a* (also known as “Peking-type” *Rhg1*), carries a distinct genetic structure. *rhg1-a* unexpectedly contains an intact and transcriptionally active retrotransposon within an intron of the key *Rhg1 α-SNAP* resistance gene in each repeat. The “Hartwig-type” SCN resistance from PI 437654 also carries the *rhg1-a* haplotype, and also carries the *RAC* retrotransposon within the *Rhg1 α-SNAP* genes.

Transposons have been coopted for the service of defense responses in both plants and animals (Tsuchiya and Eulgem, 2013; Huang et al., 2016). V(D)J recombination, which underlies the remarkable diversity of vertebrate adaptive immunity, apparently derives from a domesticated RAG-family transposase (Huang et al., 2016). *RAC* has inherent transcriptional activity and is positioned anti-sense within the first intron of the *rhg1-a α-SNAP* gene. It is unclear if *RAC* impacts *rhg1-a* function (discussed below). However, this study revealed the utility of *RAC* and the *RAC*-associated ss715606985 (G to A) SNP in more accurately identifying SCN resistance-conferring *rhg1-a* germplasm. Among the 19,645 USDA soybean accessions genotyped using the SoySNP50K iSelect BeadChip (Song et al., 2015), a few hundred accessions with a *rhg1-a*-type SNP signature apparently do not actually carry a *rhg1-a* locus. All of the *rhg1-a-*signature soybeans we examined that do not carry *RAC* encoded *rhg1-b* or *Rhg1_WT_ α-SNAP* alleles. Conversely, all examined *rhg1-a-*signature soybeans with *RAC* carried the *rhg1-a α-SNAP* allele.

Active retrotransposon families are abundant in soybean (Wawrzynski et al., 2008). *RAC* has similarities to a copia element near a *Phytophthora sojae* resistance locus identified by Bhattacharyya *et al*., however, SoyTE database searches returned no highly similar *RAC-*family TEs (Bhattacharyya et al., 1997; Du et al., 2010). Although whole-genome sequencing studies previously examined the HG type test soybean accessions, the *RAC* insertion within *α-SNAP_Rhg1_LC* was apparently omitted during the filtering steps of DNA sequence read mapping and assembly (Cook et al., 2014; Liu et al., 2017). Our findings revealed multiple *RAC*-family elements in soybean, and this abundance of *RAC-*family elements likely led to *RAC* omission from previous *rhg1-a* sequence assemblies. It is intriguing that *RAC*, at least from PI 89772, has inherent transcriptional activity and an intact ORF encoding conserved functional motifs – features of an autonomous element. Additionally, *RAC*’s near perfect identity with the Chr10 element supports that *RAC* family retrotransposons were recently active in soybean.

Host silencing of transposable elements can establish *cis-*regulatory networks where the expression of nearby host genes may also be impacted (Lisch and Bennetzen, 2011; McCue et al., 2012; McCue and Slotkin, 2012). In some cases, biotic stresses can influence transposon methylation, and subsequently, alter the expression of host genes near transposons (Dowen et al., 2012). We noted DNA methylation at the *α-SNAP*-*RAC* junctions. However, we also found evidence that *RAC* is transcriptionally active. In addition, the *Rhg1 α-SNAP* gene that contains *RAC* successfully expresses the α-SNAP*_Rhg1_*LC protein during SCN-resistance similarly to that observed for α-SNAP*_Rhg1_*HC. Future analyses of small RNAs may provide evidence of differential silencing of *RAC* or *α-SNAP_Rhg1_LC*. Cyst nematode infection of *Arabidopsis* has been reported to trigger the hypomethyation and activation of certain transposable elements, and moreover, many of these transposable elements reside near host genes whose expression is altered during syncytium establishment (Hewezi et al., 2017; Piya et al., 2017). Thus, it remains an intriguing hypothesis that *RAC* may influence the epigenetic landscape of *α-SNAP_Rhg1_LC* or the overall ∼36 kb *rhg1-a* repeat during infection, particular stresses, developmental stages or in specific tissues. Moreover, small RNAs deriving from the other *RAC*-like elements in the soybean genome could modulate *α-SNAP_Rhg1_LC* expression *in trans* (Slotkin and Martienssen, 2007; McCue et al., 2012; McCue et al., 2013). Our BLAST searches revealed at least two other *RAC*-like elements positioned intronically or adjacent to putative host defense and/or developmental genes. Future studies may interrogate if *RAC*, and/or other endogenous retrotransposons, impact the regulation of host defense gene networks in soybean.

The currently available picture of *Rhg1* haplotype evolution is incomplete and has been dominated by study of lines that are the product of ongoing selection for the most effective SCN-resistance (*e.g.* modern 10-copy *rhg1-b* and 3-copy *rhg1-a* haplotypes). The finding of *RAC* in all copies of the *rhg1-a* repeat, and in all confirmed *rhg1-a* haplotypes that were tested to date, suggests but does not confirm that *RAC* plays an adaptive role in those haplotypes. The sequence of the *Rhg1* repeat junction is identical between *rhg1-b* and *rhg1-a* haplotypes, as are many SNPs not present in the Williams 82 soybean reference genome, providing evidence of the shared evolutionary origin of *rhg1-b* and *rhg1-a* (Cook et al., 2012). The finding to date of *RAC* only in *rhg1-a* haplotypes suggests that this retroelement probably inserted in *Rhg1* after the divergence of *rhg1-b* and *rhg1-a*. However, it is also possible that the *RAC* retroelement was ancestrally present but then purged from the progenitors of current *rhg1-b* accessions. A correlation has been demonstrated between *Rhg1* copy number and SCN-resistance, and, *rhg1-a* in the absence of *Rhg4* confers only partial SCN resistance (Liu et al., 2012; Cook et al., 2014; Lee et al., 2016; Yu et al., 2016; Kandoth et al., 2017). Yet there are no known instances of *rhg1-a* accessions with an *Rhg1* copy number above three. The *RAC* may allow increased and/or more tightly regulated expression of *rhg1-a*. Alternatively, it is possible that absence of *RAC* is advantageous in allowing increased copy number of *Rhg1*, but that too is only a hypothesis, raised by the present work and in need of future testing. Additional questions about *Rhg1* locus evolution remain that have functional implications for the efficacy of SCN resistance. For example, we know of no *rhg1-a* haplotypes that carry an α-SNAP*_Rhg1_*WT-encoding *Rhg1* repeat, which all examined *rhg1-b* haplotypes do contain. Might *RAC* acquisition have influenced this absence of the WT *Rhg1* repeat? Do any α-SNAP*_Rhg1_*LC-expressing accessions exist that do not carry the *RAC* integration? The USDA soybean collection contains numerous accessions that are positive for *NSF_RAN07_* but which carry *Rhg1* copy numbers below 3 or 10, or have slight deviations from consensus *rhg1-a* or *rhg1-b* SNP signatures. Intensive study of these accessions may shed further light on *Rhg1* haplotype evolution, and moreover, may facilitate the discovery of new and agriculturally useful *Rhg1* alleles.

The finding that popular *rhg1-a* breeding sources contain an intact retrotransposon within *α-SNAP_Rhg1_LC* was surprising, given that these accessions have previously been sequenced multiple times by different groups (Cook et al., 2014; Liu et al., 2017; Patil et al., 2019). The *RAC*-SNP is rare among all USDA accessions, and approximately 700 of 19,645 USDA accessions carry a SoySNP50K signature predictive of an *rhg1-a* haplotype. However, ∼400 of these putative *rhg1-a* accessions do not carry the *RAC*-SNP and all of the non-*RAC* putative *rhg1-a* accessions that we sampled did not encode *α-SNAP_Rhg1_LC*, indicating that they are not true *rhg1-a*. Accordingly, most of these non-*RAC* accessions scored phenotypically as SCN-susceptible. Taken together with our PCR assays showing perfect correlation of the *RAC*-SNP with *RAC* presence, our findings indicate that the *RAC-*SNP successfully identifies true *rhg1-a* loci (*i.e*., those that encode the α-SNAP*_Rhg1_*LC protein) which, in combination with *Rhg4* and other loci, confers strong SCN resistance.

Correlation of SNP data with previously published SCN resistance phenotype data indicated that the vast majority of “R” scoring *rhg1-a*-signature accessions were *RAC* ^+^. Many of the 705 accessions postulated (using the earlier SoySNP50K SNP signature) to be lines that carry *rhg1-a* turned out to carry *rhg1-b*, which would explain their resistance to SCN. The large majority of the subset that are not positive for *RAC* were scored as SCN-susceptible or moderately susceptible. Some *RAC* ^+^ lines also were scored as SCN-susceptible or moderately susceptible, but most of these are apparently due to the absence of a resistance-associated *Rhg4* and/or the Chr 11 *α-SNAP* intron-retention allele (*α-SNAP_Ch11_-IR*), consistent with the established contributions of those loci to SCN resistance (Meksem et al., 2001; Yu et al., 2016; Kandoth et al., 2017). Among the “R” scoring *RAC^+^* accessions, SNPs genetically linked to *Rhg4* and the Chr 11 *α-SNAP* intron-retention allele were substantially elevated.

Potential modulation of *α-SNAP_Rhg1_LC* expression by *RAC*, either during the SCN-resistance response or in certain developmental or stress situations, could benefit *rhg1-a-*containing soybeans that have significantly depleted WT α-SNAP proteins. α-SNAPs, together with NSF, carry out essential eukaryotic housekeeping functions by maintaining SNARE proteins for vesicle trafficking. Notably, among true *rhg1-a* accessions, the abundance of wild-type (WT) α-SNAP proteins is sharply diminished, as compared to *rhg1-b*- or SCN-susceptible soybeans (Bayless et al., 2018). Moreover, α-SNAP*_Rhg1_*LC protein was shown to be cytotoxic in *Nicotiana benthamiana* while WT α-SNAP co-expression alleviated this toxicity (Bayless et al., 2016). The more recent discovery of the *NSF_RAN07_* allele as a requisite for the viability of *Rhg1* soybeans further underscores the necessity of the SNARE-recycling machinery for overall plant health (Bayless et al., 2018). The intronic copia element within the *Arabidopsis RPP7* (*Recognition of Peronospora Parasitica 7*) gene serves as an example of a retroelement with immunomodulatory function, having been shown to modulate *RPP7* transcript splicing and expression (Tsuchiya and Eulgem, 2013). However, we did not detect any influence of *RAC* on constitutive α-SNAP*_Rhg1_*LC protein expression from transgenes delivered to roots. We also did not detect a functional influence of *RAC* in its native *rhg1-a* haplotype context, insofar as that α-SNAP*_Rhg1_*LC protein abundance is successfully elevated in syncytia similarly to the reported syncytium elevation of the α-SNAP*_Rhg1_*HC protein of *rhg1-b* haplotypes, which do not carry *RAC* (Bayless et al., 2016). Hence although we did not find any true *rhg1-a* soybeans without *RAC* integrations, it remains possible that *RAC* integration was a neutral event that confers no host advantage or disadvantage.

SCN causes the most yield loss of any disease for U.S. soybean farmers and *rhg1-a* offers a potential solution to SCN populations that overcome commonly used *rhg1-b* resistance sources (Brucker et al., 2005; Niblack et al., 2008; T. W. Allen, 2017). Findings continue to emerge that further characterize different sources of SCN resistance, including exciting new findings regarding copy number variation at *Rhg4* (Patil et al., 2019). An attractive overall hypothesis for future study of *RAC* is that, in the presence of SCN or other stresses, *RAC* provides an additional regulatory layer to optimize the SCN resistance response mediated by *rhg1-a* and *Rhg4*, and/or promotes plant health in the absence of SCN. By revealing the existence of *RAC* within the important *rhg1-a* haplotype, the present study provides a marker for finding such soybeans, and expands our knowledge regarding the genetic structure and divergence of the agriculturally valuable *Rhg1* source of SCN resistance.

## Acknowledgements

This work was supported by USDA-NIFA-AFRI award 2014-67013-21775 and United Soybean Board award 1920-172-0122-B to A.F.B. The work was also supported by the National Science Foundation Graduate Research Fellowship under Grant No. (DGE-1256259) to A.M.B. We thank anonymous reviewers for their comments on a previous version of this manuscript (submitted for review December 2018).

## Author Contributions

A.M.B., R.W.Z., S.H. and A.F.B designed the research; A.M.B., R.W.Z., S.H., D.J.G. and K.K.A. performed the research; all authors analyzed data and contributed to writing the paper that was drafted primarily by A.M.B.

