## Supplemental Information for "The *rhg1-a* (*Rhg1* low-copy) nematode resistance source harbors a copia-family retrotransposon within the *Rhg1-*encoded α-SNAP gene"

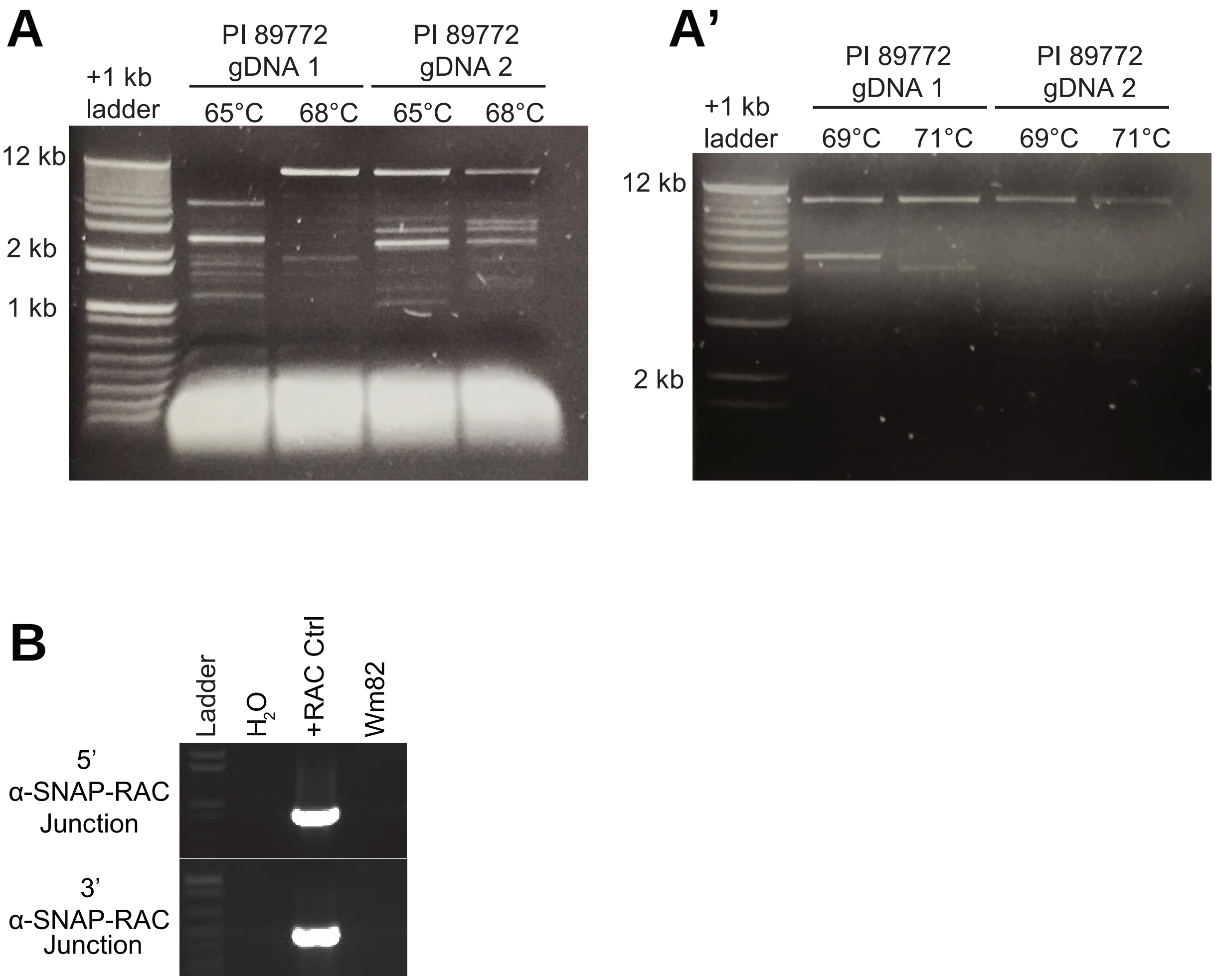


Fig. S1. Native genomic *rhg1-a* α-SNAP amplicons from PI 89772 are unexpectedly large and contain an inserted DNA element. (*A* and *A’*) Agarose gel showing PCR amplification of the native genomic *rhg1-a* α-SNAP amplicon. Similar results obtained from two independent genomic (gDNA) DNA preparations of PI 89772 and with varying primer annealing temperatures. (*B*) Agarose gel showing no detection of *RAC-α-SNAP* junctions within the WT *Rhg1* α-SNAP encoded by the soybean reference genome, Williams 82 (Wm82), relative to a positive control plasmid containing the subcloned PI 89772 (*rhg1-a*) *RAC-α-SNAP* (+RAC Ctrl); $H_2O$ - no template negative control.

#### α-SNAP$_{Rhg1}$*LC RAC* (*rhg1-a* associated copia) Nucleotide Sequence

**Features:**

*α-SNAP$_{Rhg1}$LC* Exon 1
*α-SNAP$_{Rhg1}$LC* Intron 1
*RAC* 3' LTR
*RAC* Polyprotein ORF (anti-sense to *α-SNAP$_{Rhg1}$LC* ORF)
*RAC* 5' LTR
*α-SNAP$_{Rhg1}$LC* Exon 2
Primer binding site
Putative polypurine tract (3'-GAGGGGGG-5')

ATGGCCGATCAGTTATCGAAGGGAGAGGAATTCGAGAAAAAGGCTGAGAAGAAGC
TCAGCGGTTGGGGCTTGTTTGGCTCCAAGTATGAAGATGCCGCCGATCTCTTCGAT
AAAGCCGCCAATTGCTTCAAGCTCGCCAAATCATGTTTTTCCTCTTTCTCTCTACTT
TTTTTAAATTCCATTTCGTGTCTCCTCAAAATGTTGATTTAGTGTCATAAATCATAATT
ATTATTCTCTTCTATTGTTGTTATTTTATTGTTATTACTTCAATCGACGAGTGTGTTGA
GTTTTGAGGTGTCCGATTTCCCGATTAATTGAAGTATAGTTTTAATCTGATTTTACTG
GAAAATATTTTTTTGCCTGATTTTTGTTTTTTGGAACAATTACTAGCATATAAATTAGA
ATTGTGGATGAAGTATAAGAAAACTACAGGAGCATAAGGAAGAAGAAGTGAGCTTG
AATATTTCAGAGAAGAAGATCAGCTTCAGCTACTTATTTTCGTATTAACAGAGAAGG
TTTATATACATGATGTGTGATTGTTATAACAGAAAAGCTAACTAACTCAACTAACCC
AACTACCCTTAACTGATACTGTTATACTGCTAAGAGCCCCCCTCAAGCTGGGAATG
GATATTCATCATTCCCAGCTTGTTACAGAGATGCCGAAAAACAGCAGGTGCAAGAG
CTTTGGTGAATATATCCGCTAGTTGTAAAGCTGACGAAACCGGAAGAAGCTTTAGG
AGACCTGAGTTAAGCTTTTGACGAACAATATGACAGTCTATCTCGATATGTTTAGTT
CGTTCGTGAAAAACGGGATTAGTAGCTATTTGGATGGCTGACTGATTATCACAGTA
TAAGTTCGCTGGTTGAACGAATGAGATACGAAAGTCTTGAAGCAAGAAGGTCAGCC
ACTGTAGTTCGCAAGTAGTGGAGGCAAGAGCGCGGTATTCAGCTTCGGAAGAACT
GCGTGACACAGTTGGCTGCTTCTTGGATTGCCAAGAAACCAGAGAGGAACCAAAG
TAAACTAAGTAGCCGGTAGTGGATTTCCTTGAATCTTTGCATCCAGCCCAATCCGA
GTCACTAAAGGCTCGGAGTTGTGCGGTACCTGCGGCAGTGAAGAAGATACCTGAT
CCCGGAGAACTCTTGAGGTATCGAAGAATCCGAAAGGCGGCTTGAAGATGAGCAT
TGGTGGGGGCGGCCATGTACTGGCTGAGTTGTTGAACAGCATACGTTATATCGGG
CCTGGTGTTGGTAAGGTATATTAATTTACCGATCAATCGCCGATAAGAGGAAGAAG
ACTCAGCTGAGAGAGGACTGCCCGAATCTGCCTGTAACTTCGTAGAGTAGTCTATT
GGTGTTGAATTGGGCTTGCATCCCAGCATTCCGGATGCATTTAGAATGTCTAATGT
ATACTTGCGTTGGCATAAGTGTATCCCTTTCGAGCTTCGGGCGATTTCAAGCCCAA
GGAAAAACTTTAAATCCCCAAGATCCTTGATCTTAAATTCAGAATCTAAGAGGGTGA
CAATTGTTTGTATTTCGGTCATGCTATTTCCTGTGAGAATGATGTCGTCTACATAAA
CAAGAAGGATGGTTGTGATGTTTCCAGTAAACCGCAAAAAGAGAGAGTGATCCGC

AGTTGACTGATGAAAGCCATGAGAGGTTAAGAAGCTTGACAATTTTACGAACCATT
GTCTGCTGGCTTGTTTGAGACCATAAAGAGACTTTTGAAGGCGACATACAAGCTTT
GGGTTATCAACGGAAAGTCCCGGAGGTATTTGCATATAAACCTCCTCGTCAAGTTC
TCCATGGAGGAATGCATTATTAACATCTAGCTGCCGTAAATGCCACTGGTTTAATG
CTGCAATTGCAAGAAGAAGGCGCACCGTGGTCAGCTTTGCTACCGGAGAGAAAGT
GTCAAGGTAGTCTAACCCTTCCATTTGGGTGTATCCCTTTGCAACCAGCCGCGCTT
TATGCCTTTCGATGGATCCATCTGCTCTATACTTTATTTTATAGACCCATCTGCACC
CAATAGCCGTCTTGTGAGAAGGGAGAGGTGTGAGGCGCCATGTTTGGTTCGACTG
AAGAGCTCGTAGCTCGGCTTCCATGGCCTTAATCCAGCAATCATGGCGAGAAGCA
TCGACATATGAGGTTGGCTCTGTGACGGAGGAAATATTCATGACAAAGTTCCTGTG
GGCAGGAGACAAGCGTGAGTAAGAGAGTACGGAACTAAGTGGATAACGAACAGCC
ATTGAAGTGCTTGGTGTAGAGGAAGCAAACTCTCTGTGGTAATCTCGGAGGTACGT
TGGGGTGTTTTTGGTTCTGGTGGATCGTCTAAGGTGTGAAGGAGGATGATTATGTT
CAGGTTCATTTGATGATGGTATAGAGATCATAGGTGGTGATGAAAGACTCTCTGTTT
GTGGGTCATCGTTTCTGTCGGAGAAGGATTCCGGTGAGGGAGCTGGATATTCTAA
GTGTGTATGCTGAGTTTCAGAAAGATAAGGAAAATGATCCTCATAAAATGTGATATT
TCGAGAGATGCTAACATCATTAGAGTGCAAATCATACACAAGATATCCCTTTGTATG
CATTTTAAAACCGATGAATATGCATGGATGAGCCTTAGCATCAAGCTTTTGCCGGTT
TGCCTTGAGTGTATTTATGTAACATAGACACCCGAAAACACGAAGGTTAGAAATGT
CACAAGGGTGTTTATGCAGCTTTTCATAGGGTGAAACATTATGCAAAAACGGCGTG
GGAATACAATTAATCAAGTAAGTGGCATGCGGCAAAGCGTAACACCAGAAGCTTG
GTGGTAGACTTGCCTGAAACAAAAGTGCACGTGTGACATTGAGAAGGTGCTGGTG
TTTGCGTTCTACAATTCCGTTTTGTTCTGGAGTTTCAATGCACGTGGTCTGGTGTAT
GATGCCCTTTGATGCATAGTAATGATGCATGGAGAATTCAATGCCATTATCACTTCT
GATGATCTTAACCTTGCCATCGTATTGTGTTTCAATGAATGTAATGAAGTTCATGAT
TATATGTCGGGTTTCAGCTTTGGATTTCATAAGATGAACCCATGTAAAGCGTGAGC
AATCATCAACTATAGTTAAGAAGTATTTGTGCCCATGCATGGATGGTTTAGAGCAC
GGACCCCATATGTCCATATGCAGTAAGTCAAAAATGTGAGATGCATGTGAATGGCT
AAGAGAAAAAGGTAATTTCTTTTGTTTCGCATGATGACACGTGTTGCAAACAAAATC
CTTATTATTTTTGAGAAGGGGATAGTAAGCTTTCATACATTGTATTCTTTCAGTGGAT
GGGTGGCCTAACCTAAAATGCCAAAGGTCAATAGGTATTACATTACATCGAGGGTG
AGTAATAGTGGAGTTTACGGTTTTGGTGGTCAGCTGAGCAGGTATTAAATGGTAGA
GACCGTGTTTTGCTTCAACTATACCAATCCTCATATGGCTGTTCACTTCCTGTAATA
CACACGATGTAGAGGAGAATATCAATTCACAATTAACGGAAGACACAAGTTTTGAT
ATTGAAATGATATTGAACGTAAAGGAAGGAATGTAAAGAACGTCTTGTAAAATGATG
TTTGATGAAAGCTTGACAATTCCTGAGTGGGTTGCACAAACACACTGGCCATTCGG
GAGTTTCACCGTGATAGGATCGATTTGTTTATAGGAATGAAGGTTGCGTAGGGAGT
AAGTCGCGTGGTCCGTTGCTCCTGAATCCAATATCCAGGAGACGCATGGTGTTCT
GAGAGAAAGGGACATACCTGGATTTGTCGGAGTATTGATACATGATGAGATTGAAG
CCATTTGTTTGGATTGAGAGATTGTCGTGTTTCCTGCGGATGGTTCCTGGATTAAG
GCTAGGAGTGCCTTGTACTGCTCGGGGGAGAAACGAACAGAATCATGAGCCTCGT
GGTGCTGAGCTTGATCTTCGTTAGCTTTGTTTTCAACGGCTACGAGATTGTTTACAG
TGGTTCTTCCTCCATAGGGCTTGTAACCCGGCGTGTAACCCGGTGGGTACCCGTG

TTTTCGATAACACACGTCCACAGTGTGTCCCATCTTGCCACAATGAGTGCAAGCTT
TCCTTCCACTATTCTTGTTTCTAGTATCGTGATTAGAGGGCATTCCATGTTTCTTATA
ACATGTGCTTTCTAAGTGACCAACACGTCCACAGAAGTCGCAGACAGTTTTAGCAG
CGTTTATAGAAATTTCTTTGGGTTCGAAATGAATACCTGGTCCAGCGTTTCCCAGTA
ATTGCCTTTCCTGTTGAGCCACATAGGAGAAAATCTTGGAGATAGCAGGTATAGGA
TCCATGAGAAGAACGTGGGATCGTATATTTCCGTACTGTTCATTCAGACCGCGTAG
AAACTGCATGGCCCTATCTTCGAGTTTCCGCTGTGCGATAATGGTGAATGCATTGC
AGGAACATCTTATATTACATGAGCAAATGGGATCGGGTCTAAAGTTCTCGATTTCGT
CCCAAATGACGCGTAATCGTGTGAAATACTCAGTTACTGTGAGCGTACCTTGCTTC
ATCGTCGAAGCTTCTTGTTGAAGGTCGGATATGCGTAAAAGATCTCCCTGAGAGTA
TCTTGATTTCAGATCGCGCCAAATTTCCTCGGCTTTGTCCATCCAAAGTATGCTCTG
ACGTATGGAGATGGCCACCGAATGAACTATCCACGAGACGACCATATTGTTACATC
TACGCCATGCTCCGTGCATTCTGTCCGTTTTTAGAGGTTCCGGGGCGCTGCCATCT
ATGAACTCCACTTTATTCTTGGCACTCAATGCAGTGACCATAGACCTGCTCCATGA
GTGGTAATTGGTTGAGTCTAGGACTGGGGAAACAAGAGCGATAGCTGGGTTTTCG
CTTGGATGGAGGTAGAGATAACTCTCCATGTTACTAGCAGAAGATTCGTTCATGGT
GGATAATGGAAGAATGCGCAGCAGAACTCTTCTTTTAGAAGAGCTCTGATACCATA
AGAAAACTACAGGAGCATAAGGAAGAAGTGAGCTTGAATATTTCAGAGAAGAA
GATCAGCTTCAGCTACTTATTTTCGTATTAACAGAGAAGGTTTATATACATGATGTG
TGATTGTTATAACAGAAAAGCTAACTAACTCAACTAACCCAACTACCCTTAACTGAT
ACTGTTATACTGCTAAGAAAGTACGACAATCAACTCTGTGTTGTTTGTGACTACGCT
CACTTTCAATTTGACGACTAATCTCTTTATTTTGTTGAAAGTGACGAACTTTGAAATT
GATGTTGGAATAGTTCTGTTTATTGTTCTTGATTTGATCTATGTGGCATTTTAGGGG
ACAAGGCTGGAGCGACATACCTGAAGTTGGCAAGTTGTCATTTGAAG

Fig S2. Complete nucleotide sequence of the PI 89772 (*rhg1-a*, *Rhg1* low-copy) *RAC* (element and flanking exonic *α-SNAP*$_{Rhg1}$*LC* regions. Key sequence features are color coded as indicated in the above box.

#### S3 PI 89772 *rhg1-a RAC*-encoded polyprotein (1438 residues)

MESYLYLHPSENPAIALVSPVLDSTNYHSWSRSMVTALSAKNKVEFIDGSAPEPLKTDR
MHGAWRRCNNMVVSWIVHSVAISIRQSILWMDKAEEIWRDLKSRYSQGDLLRISDLQQ
EASTMKQGTLTVTEYFTRLRVIWDEIENFRPDPICSCNIRCSCNAFTIIAQRKLEDRAMQ
FLRGLNEQYGNIRSHVLLMDPIPAISKIFSYVAQQERQLLGNAGPGIHFEPKEISINAAKT
VCDFCGRVGHLESTCYKKHGMPSNHDTRNKNSGRKACTHCGKMGHTVDVCYRKHG
YPPGYTPGYKPYGGRTTVNNLVAVENKANEDQAQHHEAHDSVRFSPEQYKALLALIQ
EPSAGNTTISQSKQMASISSCINTPTNPGMSLSLRTPCVSWILDSGATDHATYSLRNLH
SYKQIDPITVKLPNGQCVCATHSGIVKLSSNIILQDVLYIPSFTFNIISISKLVSSVNCELIFS
STSCVLQEVNSHMRIGIVEAKHGLYHLIPAQLTTKTVNSTITHPRCNVIPIDLWHFRLGH
PSTERIQCMKAYYPLLKNNKDFVCNTCHHAKQKKLPFSLSHSHASHIFDLLHMDIWGP
CSKPSMHGHKYFLTIVDDCSRFTWVHLMKSKAETRHIIMNFITFIETQYDGKVKIIRSDN
GIEFSMHHYYASKGIIHQTTCIETPEQNGIVERKHQHLLNVTRALLFQASLPPSFWCYAL
PHATYLINCIPTPFLHNVSPYEKLHKHPCDISNLRVFGCLCYINTLKANRQKLDARAHPC
IFIGFKMHTKGYLVYDLHSNDVSISRNITFYEDHFPYLSETQHTHLEYPAPSPESFSDRN
DDPQTESLSSPPMISIPSSNEXEHNHPPSHLRRSTRTKNTPTYLRDYHREFASSTPSTS
MAVRYPLSSVLSYSRLSPAHRNFVMNISSVTEPTSYVDASRHDCWIKAMEAELRALQS
NQTWRLTPLPSHKTAIGCRWVYKIKYRADGSIERHKARLVAKGYTQMEGLDYLDTFSP
VAKLTTVRLLLAIAALNQWHLRQLDVNNAFLHGELDEEVYMQIPPGLSVDNPKLVCRLQ
KSLYGLKQASRQWFVKLSSFLTSHGFHQSTADHSLFLRFTGNITTILLVYVDDIILTGNS
MTEIQTIVTLLDSEFKIKDLGDLKFFLGLEIARSSKGIHLCQRKYTLDILNASGMLGCKPN
STPIDYSTKLQADSGSPLSAESSSSYRRLIGKLIYLTNTRPDITYAVQQLSQYMAAPTNA
HLQAAFRILRYLKSSPGSGIFFTAAGTAQLRAFSDSDWAGCKDSRKSTTGYLVYFGSS
LVSWQSKKQPTVSRSSSEAEYRALASTTCELQWLTFLLQDFRISFVQPANLYCDNQSA
IQIATNPVFHERTKHIEIDCHIVRQKLNSGLLKLLPVSSALQLADIFTKALAPAVFRHLCNK
LGMMNIHSQLEGGS*

##### Conserved Structural Features of Copia Polyproteins

GAG binding motif: CTHCGKMGHTVDVC

Protease motif: WILDSGATDH

Integrase (GKGY motif): GCLCYINTLKANRQKLDARAHPCIFIGFKMHTKGY

Rev. Transcriptase motif:
QLDVNNAFLHGELDEEVYMQIPPGLSVDNPKLVCRLQKSLYGLKQASRQWFVKLSSFL
TSHGFHQSTADHSLFLRFTGNITTILLVYVDDII

Fig. S3. Translation of the *RAC*-encoded polyprotein (1438 residues) from accession PI 89772. Conserved features of the polyprotein are colored and identified as indicated in the above box.

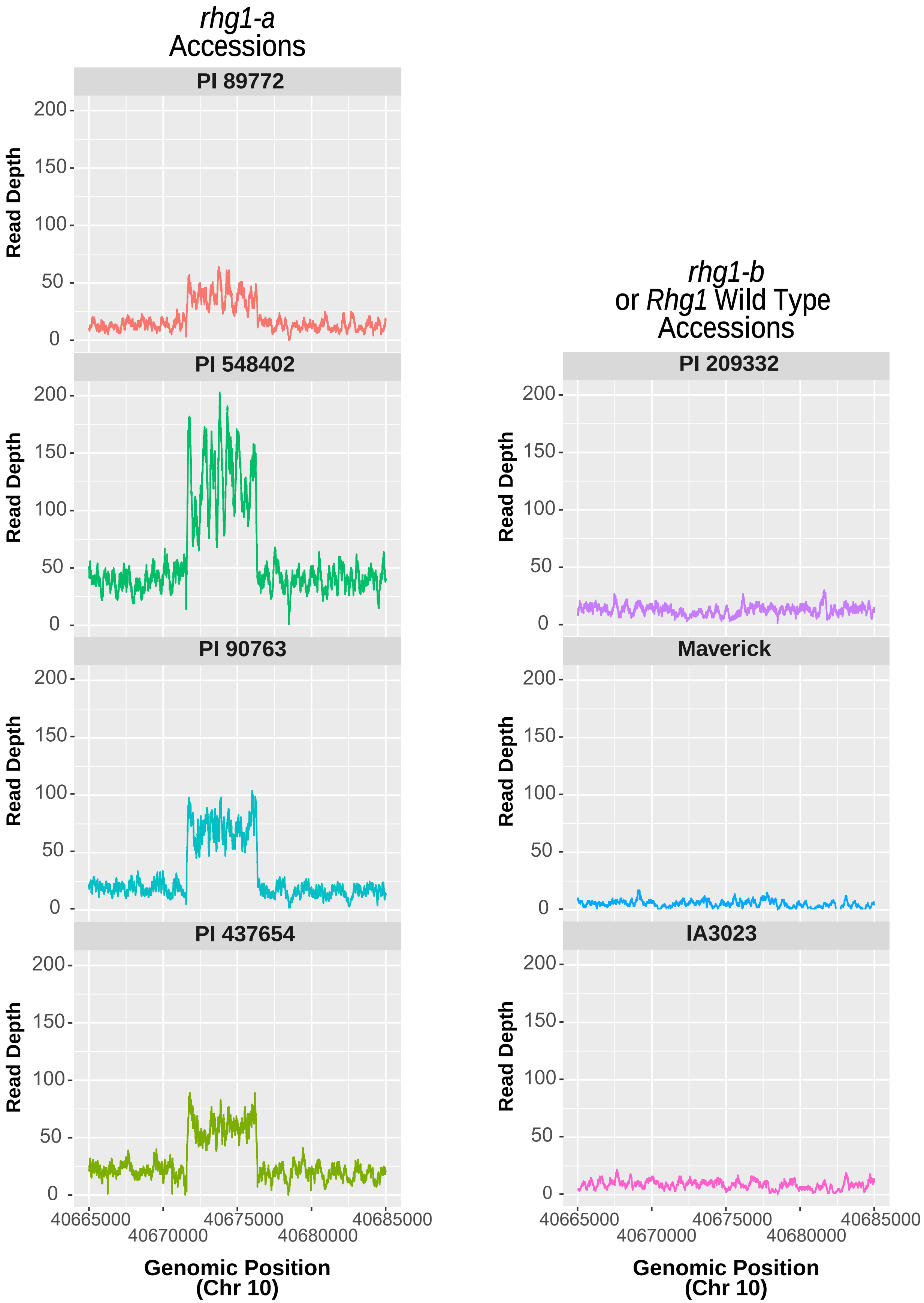


Fig. S4. Depth of short genomic reads that align to the *RAC* nucleotide sequence. The *α-SNAP-RAC* insertion is not present within the Wm 82 soybean reference genome, thus reads allign to the genomic position of the Chr 10 *RAC*-like element (99.7% nucleotide identity to *RAC*). Read depth for *rhg1-a* accessions (PI 89772, PI 548402, PI 90763, PI 437654) is ~3 to 4 fold greater than for the *rhg1-b* accessions (PI 209332, Maverick) and $Rhg1_{WT}$ accession (IA3023), which do not contain Chr 18 *Rhg1 α-SNAP-RAC* insertions.

**A**

```
RAC          AAGCACATGTTATAAGAAACATGGAATGCCCTCTAATCACGATACTAGAAACAAGAATAG
C10          AAGCACATGTTATAAGAAACATGGAATGCCCTCTAATCACGATACTAGAAACAAGAATAG
             ************************************************************

RAC          TGGAAGGAAAGCTTGCACTCATTGTGGCAAGATGGGACACACTGTGGACGTGTGTTATCG
C10          TGGAAGGAAAGCTTGCACTCATTGTGGCAAGATGGGGCACACTGTGGACGTGTGTTATCG
             ************************************ ***********************

RAC          AAAACACGGGTACCCACCGGGTTACACGCCGGGTTACAAGCCCTATGGAGGAAGAACCAC
C10          AAAACACGGGTACCCACCGGGTTACACGCCGGGTTACAAGCCCTATGGAGGAAGAACCAC
             ************************************************************
```

**B**

| SoySNP 50K | *rhg1-a* | *rhg1-b* |
|---|---|---|
| ss715629217 | C | C |
| ss715629233 | C | C |
| ss715629242 | C | C |
| ss715629248 | C | C |
| ss715629260 | G | G |
| ss715629264 | **A** | **G** |
| ss715629266 | T | T |
| ss715629273 | G | G |
| ss715629280 | G | G |
| ss715629286 | **T** | **C** |
| ss715629288 | **C** | **T** |
| ss715629291 | A | A |
| ss715629296 | C | C |
| ss715629298 | **G** | **A** |

**C**

| Accession | *RAC* SNP? | Detected *Rhg1* α-SNAP | True *rhg1-a*? |
|---|---|---|---|
| PI 417441 | No | HC α-SNAP | No |
| PI 507148 | No | WT α-SNAP | No |
| PI 458094 | No | HC α-SNAP | No |
| PI 408304 | No | HC α-SNAP | No |
| PI 603438B | No | HC α-SNAP | No |
| PI 567319B | Present | LC α-SNAP | Yes |
| PI 567234B | Present | LC α-SNAP | Yes |

Fig. S5. Sequence of the *RAC* SNP (ss715606985), nucleotide SNP signatures of consensus *rhg1-a* (*Rhg1* low-copy) and *rhg1-b* (*Rhg1* high-copy) haplotypes, and detected *Rhg1* α-SNAP transcripts from *RAC*$^{+}$ or *RAC*$^{-}$ accessions with consensus *rhg1-a* SNP-signatures. (*A*) Short DNA alignment of *RAC* and the Chr10 ("C10") element showing the position of the ss715606985 SNP (G to A) associated with *RAC* presence. (*B*) Consensus SNP signatures for *rhg1-a* and *rhg1-b* haplotypes, as identified by Lee *et al*, 2015. (*C*) *Rhg1* α-SNAP alleles detected from genomic DNA subclones of *RAC*$^{+}$ or *RAC*$^{-}$ acces-sions which have a consensus *rhg1-a* SNP signature. HC, LC or WT refers to high-copy (*rhg1-b*), low-copy (*rhg1-a*) or wild-type *Rhg1* α-SNAP alleles, respectively.

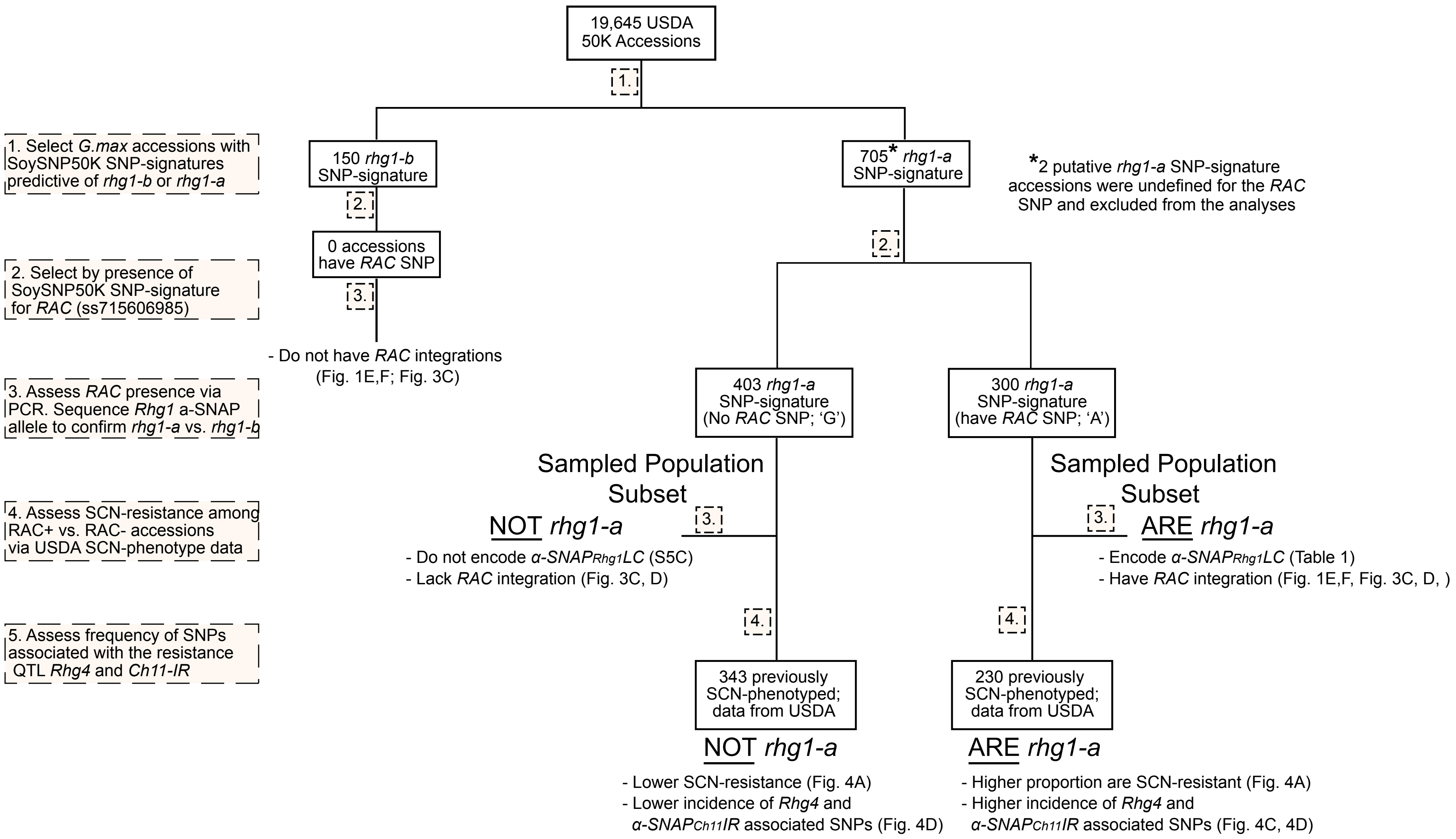


Fig. S6. Flow chart summarizing findings regarding *RAC*$^{+}$ vs. *RAC*$^{-}$ accessions, which otherwise have consensus SNP signatures for *rhg1-a*. The *RAC* SNP is useful to identify true *rhg1-a* accessions with strong SCN-resistance. See manuscript text for full description.

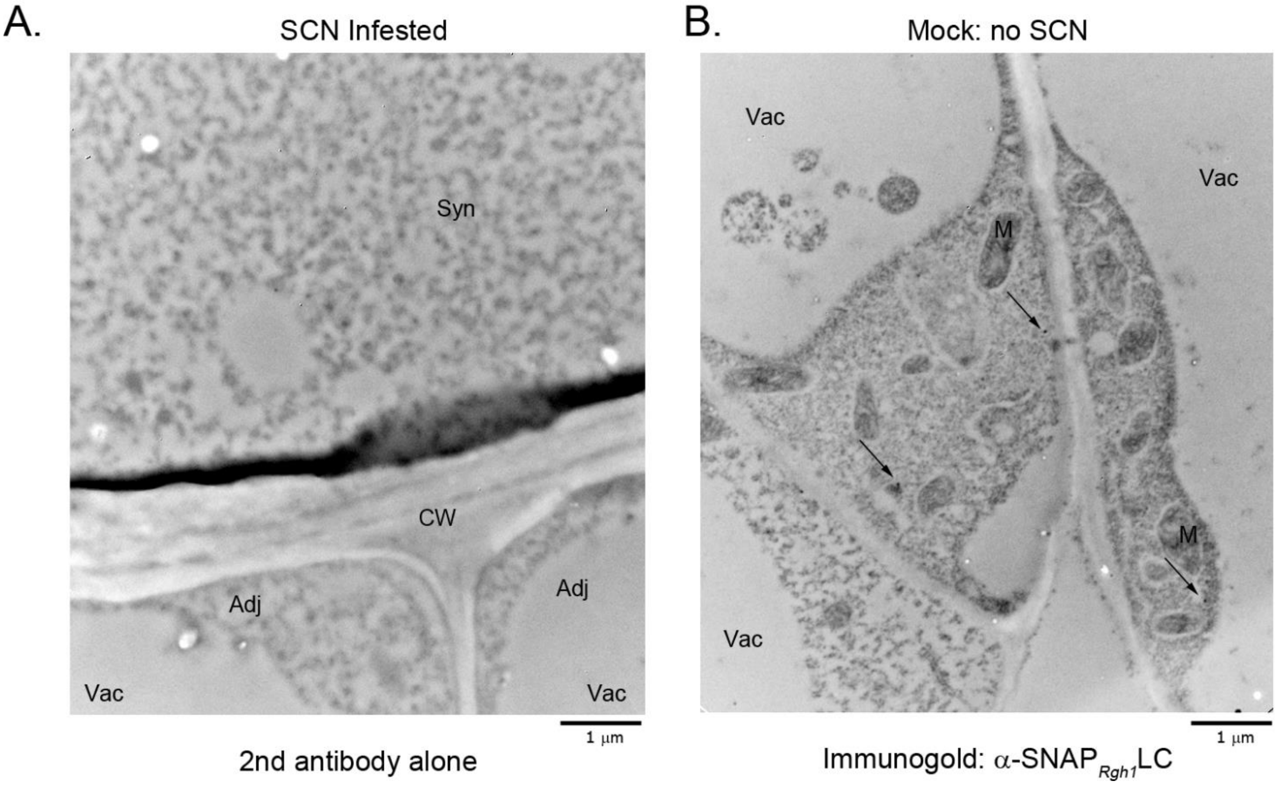


**Fig. S7** α-SNAP$_{Rhg1}$LC immunolabeling in ‘Forrest’ roots is highly specific. (A) Representative electron microscope image showing immunogold labeling using only secondary goat anti-rabbit antibody on SCN-infested ‘Forrest’ roots. Without prior incubation with the anti-α-SNAP$_{Rhg1}$LC antibody, few gold particles were observed in syncytial vs. adjacent non-infected cells from the same root sections mounted on different grids. Syn, syncytium cells; Adj, adjacent cells. (B) Electron micrograph of a mock-inoculated ‘Forrest’ root after immunogold label detection using the anti–α-SNAP$_{Rhg1}$LC primary antibody. Arrows indicate three immunogold particles across three root cell regions.

(In both A and B, CW, cell wall; M, mitochondrion; Vac, vacuole; bar = 1 µm.)

### Table S1

| Location | BLAST Score | *RAC* Identity | Note |
|---|---|---|---|
| Ch10 | 8540 | 99.70% | Copia ORF fully intact; would encode 1438 residue polyprotein |
| Ch18 | 8076 | 97.60% | Integrated within intron 1 of *Glyma.18G268000* (anti-sense, copia ORF interrupted at residue 993) |
| Ch09 | 6522 | 90% | Copia ORF adjacent and intact; anti-sense to *Glyma.09G206300* ( ABC transporter) |
| Ch14 | 4731 | 82% | copia ORF interrupted at residue 545 |
| Ch02 | 4666 | 82% | copia ORF interrupted at residue 115 |
| Ch20 | 4644 | 82% | Intronic integration within *Glyma.20G250200* (BAR domain) |
| Ch04 | 4080 | | |
| Ch01 | 3927 | | |
| Ch15 | 3910 | | |
| Ch05 | 3479 | | |
| Ch06 | 3285 | | |
| Ch03 | 3057 | | |
| Ch07 | 3036 | | |
| Ch16 | 2947 | | |
| Ch13 | 2922 | | |
| Ch08 | 2156 | | |
| Ch12 | 2040 | | |
| Ch17 | 1563 | | |
| Ch11 | 827 | | |
| Ch19 | 553 | | |
| *P.vulgaris* Ch02 | 378 | | |

Table S1. *RAC*-like elements identified from NBLAST searches of *RAC* against the Williams 82 reference genome at Phytozome.org. The highest *RAC*-like match identified in the genome of the common bean (*Phaseolus vulgaris*) is also included. Only a single element (with highest identity to *RAC*) is shown for each of the n = 20 chromosomes of soybean. Several *RAC*-subfamily elements within the soybean reference genome are also inserted into host genes, and/or include elements with intact ORFs.
